# RNA polymerase II clusters form in line with liquid phase wetting of chromatin

**DOI:** 10.1101/2021.02.03.429626

**Authors:** Agnieszka Pancholi, Tim Klingberg, Weichun Zhang, Roshan Prizak, Irina Mamontova, Amra Noa, Andrei Yu Kobitski, Gerd Ulrich Nienhaus, Vasily Zaburdaev, Lennart Hilbert

## Abstract

It is essential for cells to control which genes are transcribed into RNA. In eukaryotes, two major control points are recruitment of RNA polymerase II (Pol II) into a paused state and subsequent pause release to begin transcript elongation. Pol II associates with macromolecular clusters during recruitment, but it remains unclear how Pol II recruitment and pause release might affect these clusters. Here, we show that clusters exhibit morphologies that are in line with wetting of chromatin by a liquid phase enriched in recruited Pol II. Applying instantaneous structured illumination microscopy and stimulated emission double depletion microscopy to pluripotent zebrafish embryos, we find recruited Pol II associated with large clusters, and elongating Pol II with dispersed clusters. A lattice kinetic Monte Carlo model representing recruited Pol II as a liquid phase reproduced the observed cluster morphologies. In this model, chromatin is a copolymer chain containing regions that attract or repel recruited Pol II, supporting droplet formation by wetting or droplet dispersal, respectively.

## Introduction

Eukaryotic cells have an extensive library of genetic DNA sequences at their disposal, but selectively transcribe only a small subset of this genetic information into RNA transcripts at any given point in time. For the vast majority of genes whose transcription is controlled, the synthesis of RNA transcripts is carried out by the multi-protein complex RNA polymerase II (Pol II). Two major points at which transcription by Pol II is controlled are recruitment and pause release^1^. Recruitment brings Pol II into the vicinity of the promoter region, a sequence located upstream of an actual gene, which integrates many regulatory influences (Fig. 1A). Some of the recruited Pol II complexes engage with DNA at the promoter and start synthesizing the RNA transcript in a process called initiation. After proceeding for 20-60 base pairs along the DNA sequence, the initiated Pol II complexes enter a state of promoter-proximal pausing^2^. The rates of Pol II recruitment and subsequent release from the paused state are under cellular control and can differ between different genes and for different stimuli the cell is exposed to^1,3^. For genes with a pause release rate similar to or greater than the rate of recruitment, Pol II proceeds into proper elongation without notable retention in the promoter-proximal position^4^. In contrast, when pause release is slower than recruitment, Pol II is retained in the promoter-proximal position, thus entering a so-called poised state. Genes that exhibit poising remain ready for induction, enabling, for example, an extensive transcriptional response to heat shock^5^ and potentially the trans-differentiation of neuronal types^6^. In early embryonic development, some genes are also poised – supposedly in preparation for subsequent expression during cell type specification^4^.

**Fig. 1:**
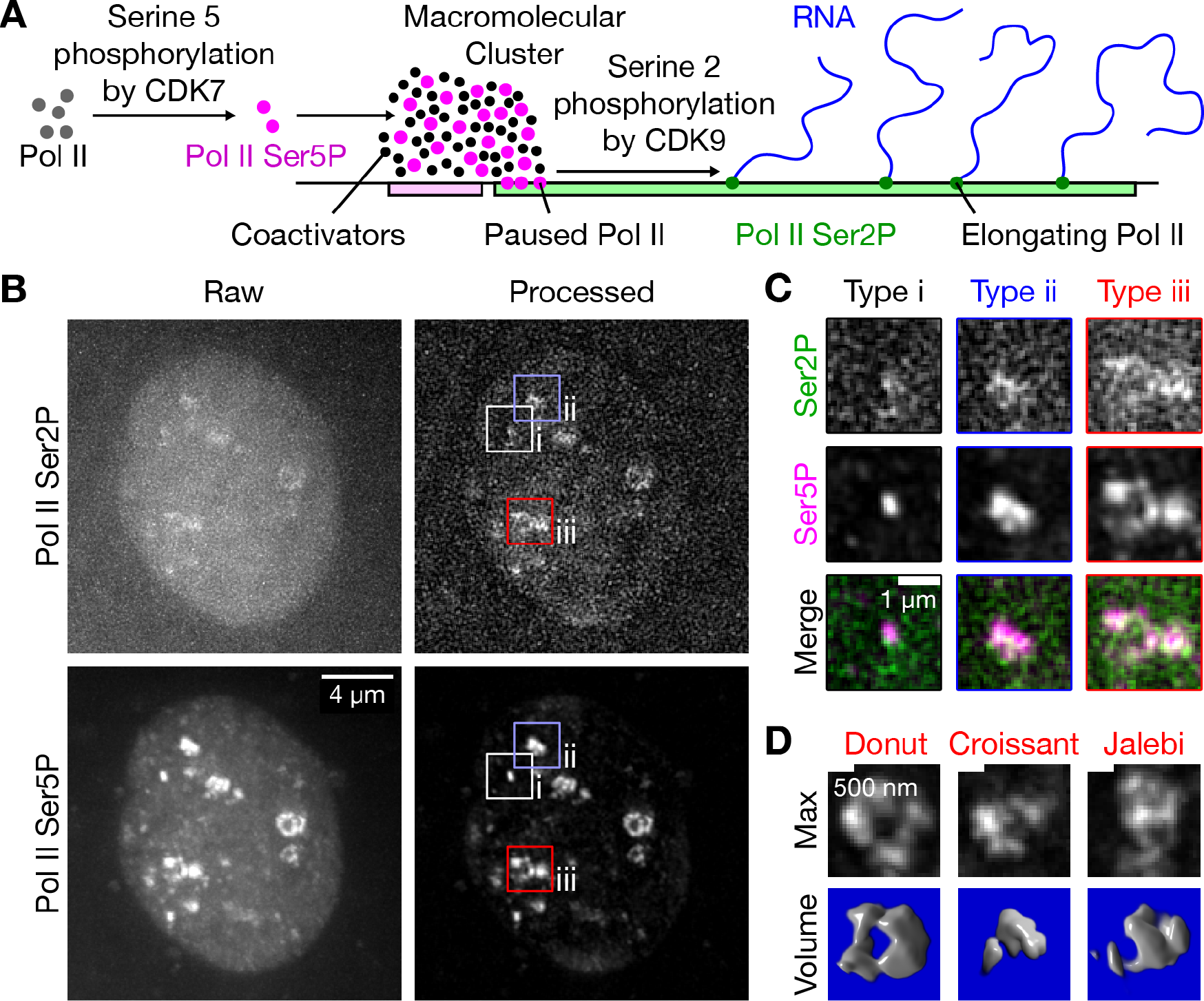
Phosphorylation-specific detection of RNA polymerase II reveals clusters displaying a variety of morphologies. A) Sketch of the recruitment and release of RNA polymerase II (Pol II) in the context of macromolecular clusters. B) Representative maximum projection of a nucleus in a live zebrafish embryo (sphere stage), where Pol II was detected via fluorescently labeled antibody fragments (Fab) specific against Ser2 and Ser5 phosphorylation (Pol II Ser2P, Pol II Ser5P). Pol II Ser5P clusters representing the different apparent types of morphologies are marked. Single time point z stacks were recorded using an instantaneous Structured Illumination Microscope (instant-SIM) microscope, raw data were processed by local background subtraction. C) Detail views of the clusters of the apparent morphological types i-iii, as marked in panel A. D) Examples of the varied morphologies of type iii clusters, shown as maximum projections and corresponding volume renderings of the processed Pol II Ser5P signal. Morphologies are named by similarity to patisserie and candy items. (3D renderings ImageJ Volume Viewer plugin).

A complementary perspective on Pol II recruitment and pause release considers changes in the localization of Pol II (Fig. 1A). It has been proposed that transcription occurs in static factories, containing several Pol II complexes and several genes in a shared local context^7,8^. This initial transcription factory picture was refined based on live-cell microscopy, showing dynamic macromolecular clusters that are enriched in Pol II and provide platforms for the initiation of transcript elongation^9–11^. These clusters are supposedly formed by liquid-liquid phase separation (LLPS) and support the co-association of activating factors such as the protein Mediator or activity-inducing chromatin remodellers^12–14^. Post-translational modifications that occur during Pol II recruitment have been found to control the association of Pol II with such clusters. Specifically, Pol II recruitment proceeds in conjunction with phosphorylation of serine 5 (Ser5P) of the Pol II carboxy-terminal domain (CTD) YSPTSPS repeat array. This Ser5P mark triggers recruitment of Pol II into clusters enriched in cyclin-dependent kinase 9 (CDK9)^15,16^. CDK9, in turn, phosphorylates serine 2 (Ser2P) of the Pol II CTD-repeat, a modification that is essential to enable pause release and subsequent elongation^17^. The newly deposited Ser2P mark also abolishes the affinity of Pol II for the Pol II-enriched clusters ^15^, allowing its relocation towards nuclear speckles associated with further RNA processing^18,19^.

Recent live-cell microscopy results suggest that the progression of Pol II through recruitment and pause release might also affect the internal organization of Pol II-enriched macromolecular clusters^20^. Such reorganization processes might be relevant to the complex morphology of Pol II-enriched clusters, which deviates markedly from the round, droplet-like shapes typical for canonical LLPS ^13,21,22^. Here, we analyze how Pol II recruitment and pause release are related to cluster morphology by a combination of live-cell and super-resolution microscopy in zebrafish embryos with lattice simulations of liquid phase condensation on block copolymers. We observe that clusters exhibit various types of morphologies that are associated with different levels of Pol II Ser5P and Pol II Ser2P. Our lattice simulations reproduce this observation, explaining cluster formation via the wetting of regulatory genomic regions by a liquid phase that is enriched for recruited Pol II, and cluster dispersal via the exclusion of elongating Pol II from this liquid phase. The causal relevance of Pol II phosphorylation is supported by the application of chemical inhibitors of Pol II phosphorylation, which induces changes in cluster morphology and cluster number that are in line with predictions from our lattice simulations. In combination with previous work on Pol II liquid phase behavior^13,14^ and studies showing condensation of transcription factors by wetting of DNA *in vitro*^23,24^, our findings in zebrafish, an embryonic model system, suggest that similar liquid phase wetting of chromatin might also occur *in vivo.*

## Results

### Recruited RNA polymerase II occurs in clusters exhibiting different types of morphologies

To study Pol II-enriched clusters, we used zebrafish embryos in the pluripotent stage of development (sphere) as an experimental model system. Zebrafish embryos provide the context of a normally developing vertebrate, and are amenable to study by light microscopy. Our previous work demonstrated that fluorescently labeled antibody fragments (Fab) against post-translational modifications are well-tolerated and provide good sensitivity as well as time resolution in zebrafish embryos^25,26^. To assess Pol II specifically in the recruited and elongating states, we injected embryos with antibody fragments against Pol II Ser5P and Pol II Ser2P, respectively. We acquired microscopy images from live embryos using an instantaneous Structured Illumination Microscope (instant-SIM), which provides approximately two-fold increased resolution in all three spatial dimensions relative to conventional confocal microscopy^27^.

In our microscopy images, the Pol II Ser5P channel (recruited Pol II) revealed distinct clusters with a rich array of morphologies (Fig. 1B). These clusters as well as their specific morphologies were long-lived, persisting for over ten minutes (Supplementary Fig. 7). The observation of long-lived clusters is in line with results from another model of pluripotency, mouse embryonic stem cells^13^. Based on the shapes of the clusters observed in the Pol II Ser5P channel, we gained the impression that clusters mostly occurred in three distinct morphological types (Fig. 1C). These morphological types seem to correlate with different levels of Pol II Ser5P and Pol II Ser2P signal (elongating Pol II). Type i clusters are small, appear as dots in the Pol II Ser5P channel, and exhibit low Pol II Ser2P signal. Type ii clusters are larger, relatively compact in the Pol II Ser5P channel, and also exhibit low Pol II Ser2P signal. Type iii clusters are also larger, appear dispersed in the Pol II Ser5P channel, and have relatively high Pol II Ser2P signal. The dispersed type iii clusters show especially complex shapes with a large morphological variety. Illustrative examples include shapes akin to donuts (a ring with a hole in the center), croissants (a crescent shape), or Jalebi sweets (segments running across each other) (Fig. 1D). Taken together, we found that recruited Pol II forms distinct, long-lived clusters with a rich array of morphologies that appear to vary with the levels of Pol II Ser2 and Ser5 phosphorylation present at a given cluster.

### Levels of recruited and elongating Pol II correlate with cluster morphology

To systematically characterize cluster morphologies and their dependence on Pol II pausing and elongation, we assessed clusters by super-resolution microscopy in fixed embryos. Specifically, we applied STimulated Emission Double Depletion (STEDD) microscopy, which significantly reduces background from low frequency contributions and out-of-focus light relative to conventional STED microscopy^28–30^. Here, we super-resolved the Pol II Ser5P distribution and acquired the level of Pol II Ser2P in a second color channel in the same focal plane by regular confocal microscopy. The Pol II Ser5P channel revealed the same apparent cluster morphologies that were seen in our live imaging data (Fig. 2A,B). The improved signal-to-noise ratio relative to our live imaging data revealed an additional detail: the elevated Pol II Ser2Phos signal associated with type iii clusters was placed directly adjacent to, but did not strongly overlap with the Pol II Ser5P signal (Fig. 2B). To more comprehensively assess cluster morphology, we extracted individual clusters and characterized their morphology by size (area) and compactness (solidity) using an automated analysis pipeline (Fig. 2C). Based on area and solidity, we gated the clusters into extreme examples of the morphology types i-iii (Fig. 2C). The gated sets of clusters exhibited systematic differences in Pol II Ser2P and Pol II Ser5P intensities: small clusters (type i) had low Pol II Ser5P and Pol II Ser2P levels; large and round clusters (type ii) had the highest levels of Pol II Ser5P and high levels of Pol II Ser2P; large and dispersed clusters (type iii) had intermediate levels of Pol II Ser5P and high levels of Pol II Ser2P (Fig. 2D). All the results obtained by the analysis of STEDD images were reproduced, at lower resolution, in an analysis of our live imaging data (Supplementary Fig. 9). Together, these results suggest a direct correlation between the morphology and phosphorylation levels of Pol II clusters (Fig. 2E).

**Fig. 2:**
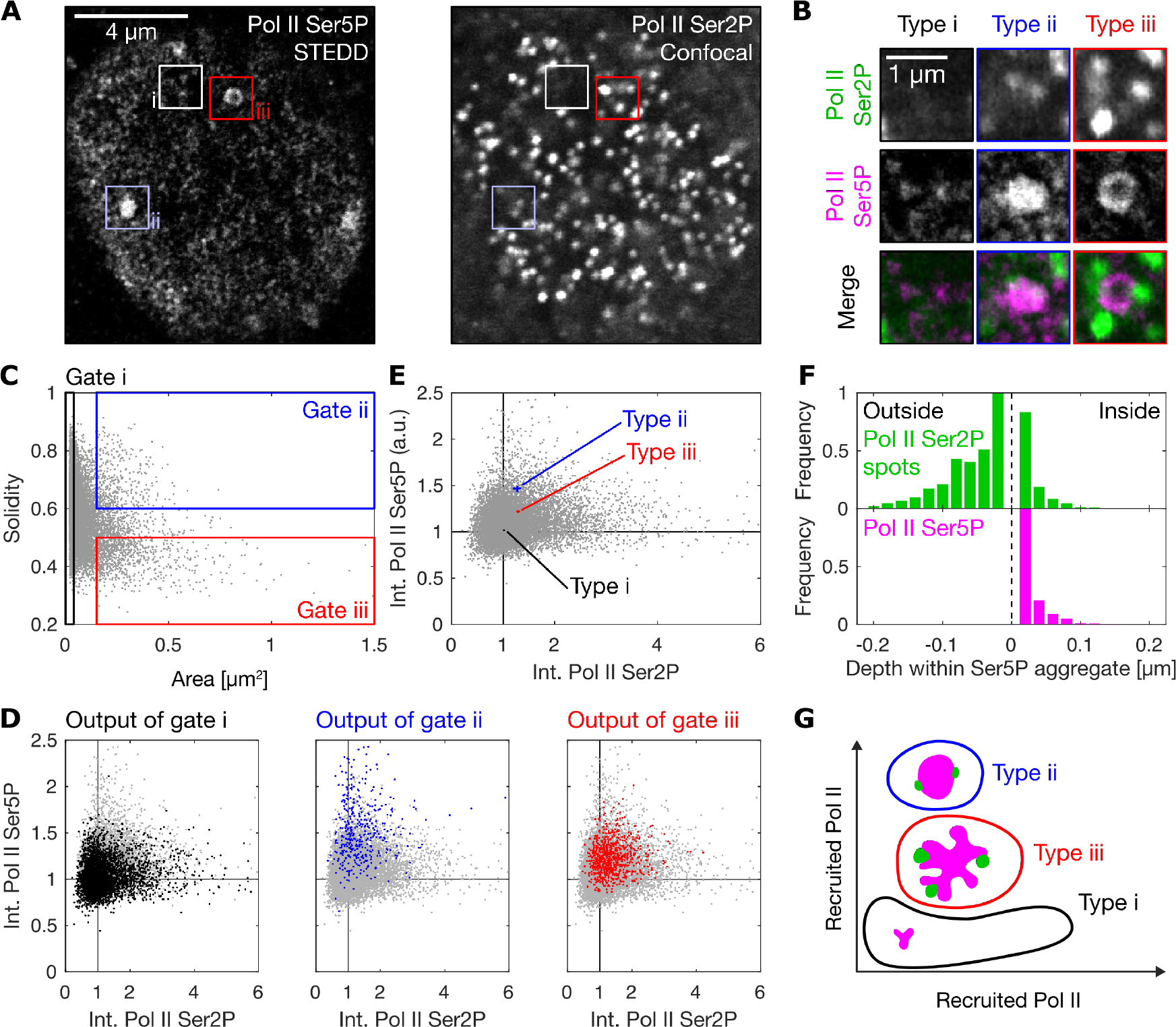
Super-resolution microscopy reveals types of cluster morphology correlating with levels of recruited and elongating RNA polymerase II. A) Representative nuclear mid-section obtained by STEDD super-resolution microscopy from a fixed late blastula stage zebrafish embryo. Pol II Ser5P intensity distributions were obtained by STEDD microscopy, Pol II Ser2P intensity distributions by regular confocal microscopy from the same focal plane. Pol II Ser5P clusters with typical morphologies i-iii are marked. B) Detail views of the marked clusters, representing the typical morphologies i-iii. C) Area and solidity of individual clusters, with gate regions for the typical morphologies i-iii. Clusters were segmented based on Pol II Ser5P intensity, data obtained from a total of 52 mid-nuclear sections from two different samples. D) The Pol II Ser5P and Pol II Ser2P intensities (mean intensity across all pixels inside a given cluster’s segmentation mass) of the clusters in the gates i-iii are plotted in color over the entire ungated cluster population (light gray). Mean Pol II Ser5P and Pol II Ser2P intensities were scaled by the median value for each nucleus, then pooled. E) The median of the Pol II Ser5P and Pol II Ser2P levels of the gated clusters in panel D is plotted over the ungated population of clusters. Each cluster type is plotted with 95% bootstrapped confidence intervals in Pol II Ser5P and Pol II Ser2P direction (10,000 resamples). F) Analysis of the placement of Pol II Ser2P spots relative to Pol II Ser5P clusters (spots segmented based on Pol II Ser2P channel). Histograms show the distribution of distances to the nearest surface of a Pol II Ser5P cluster (Euclidean metric, applied to individual pixels, negative values are from outside, positive values from inside a Pol II Ser5P cluster). The distance distribution of pixels inside Pol II Ser5P clusters is shown for reference. G) Sketch of apparent morphological types of Pol II Ser5P clusters, placed by their levels of elongating and paused Pol II.

The question arises whether differently phosphorylated Pol II might in fact exert a systematic effect on the morphology of Pol II clusters. Visual inspection of our STEDD micrographs suggests that Pol II Ser5P and Pol II Ser2P form intricate structures: they are separated but in close neighborhood of each other (Fig. 2B). In particular, a comprehensive analysis shows that Pol II Ser2P clusters are placed at the margins of Pol II Ser5P clusters (Fig. 2F). This arrangement is suggestive of a scenario where Pol II Ser5P is associated with a convergent force that establishes compact clusters; Pol II Ser2P with a dispersing force that pulls outward on the margins of a given cluster. The different morphological types, their respective levels of recruited and elongating Pol II, as well as their internal organization are sketched in Figure 2G.

### Cluster morphologies are reproduced by a model of droplet condensation on regulatory genomic regions

Formation of macromolecular clusters at genomic target regions has recently been described using a model of liquid phase condensation on polymers as microscopic surfaces^13,14,23,24^. To test whether such a model can reproduce the different cluster morphologies seen in our experiments, we implemented corresponding lattice kinetic Monte Carlo (LKMC) simulations^31,32^ (for details see Supplementary Material). The LKMC simulation framework has been used previously to simulate chromosome dynamics ^25,32^ and provides a coarse-grained representation of the relevant macromolecules, in line with estimates of a few hundred Pol II complexes per cluster^13^. Specifically, we introduced a “red” particle species that represents the material forming the Pol II-enriched clusters. In line with previous work^13,19^, particles of this species exhibit self-affinity (*w_Pol-Pol_* < 0) that can, given sufficiently high affinity and bulk concentration, support formation of a concentrated droplet phase (Fig. 3A). Previous work suggested, however, that the formation of Pol II-enriched clusters does not proceed by canonical LLPS, but rather occurs by condensation specifically on regulatory sites within chromatin^13,14,22^. We thus introduced linear block polymers representing chromatin. These block copolymers contain “blue” subregions that represent extensive regulatory regions (referred to as PC, for poised chromatin) and have an affinity to the red particles (*w_PC-Pol_* < 0, see Fig. 4B)^6^. The linear polymers also contain neutral “black” segments, which have no affinity to red particles (referred to as IC, for inactive chromatin). The addition of such a polymer can facilitate cluster formation even under conditions where red particles would otherwise not phase-separate (Fig. 3A,B, for adjustment of the concentration of red particles and *w_PC-Pol_* see Supplementary Fig. 11). This behavior is typical of liquid phase condensation on a surface, also called wetting^24^. To confirm that the simulations exhibit surface-mediated condensation, we demonstrate another key behavior: the size of clusters is controlled by the amount of available surface (Fig. 3C)^24^.

**Fig. 3:**
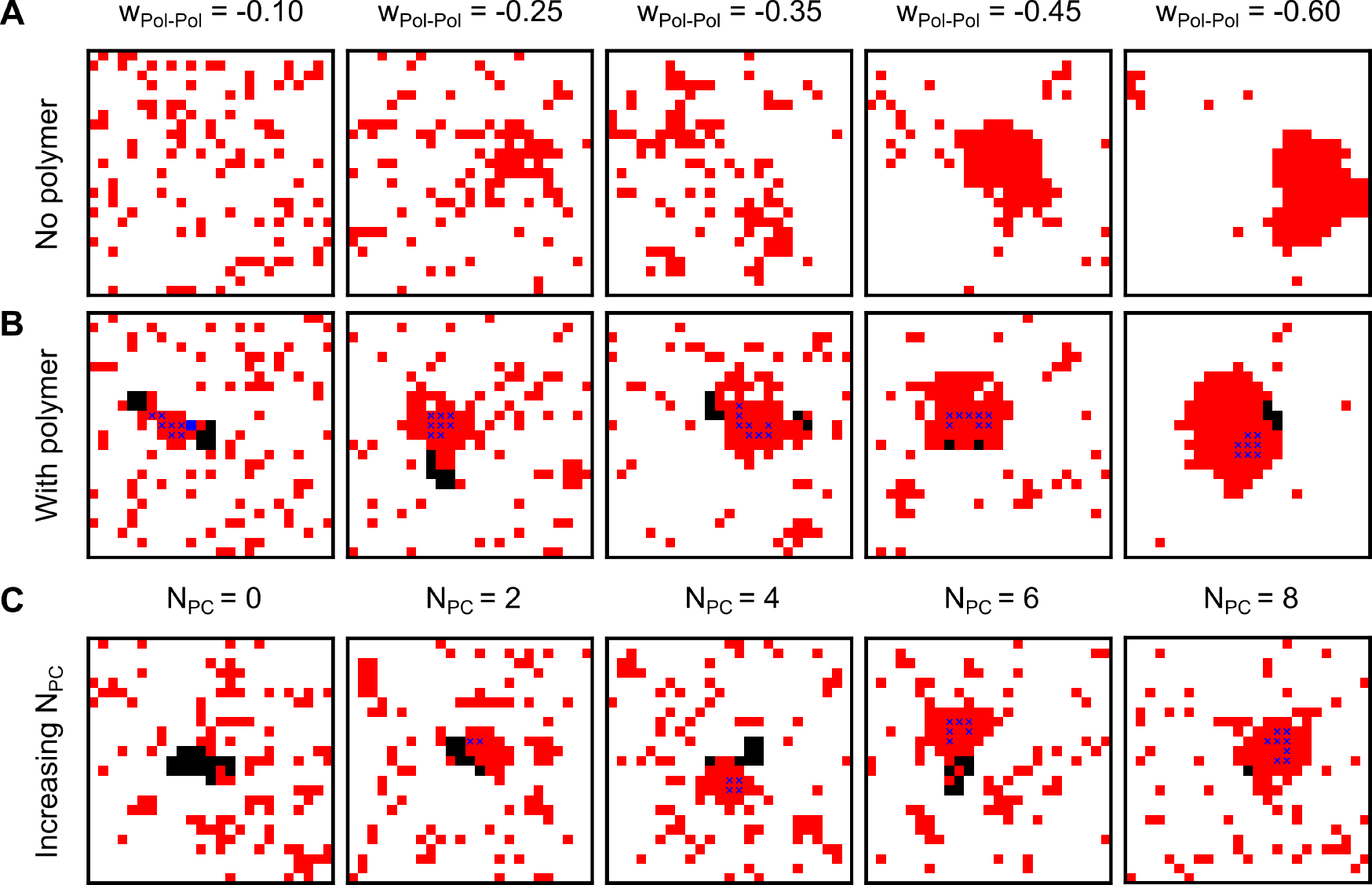
A lattice model exhibits key characteristics of liquid phase condensation with a polymeric subregion as a surface. A) Examples of lattice configurations obtained from simulations containing only red particles with increasing self-affinity (*w_Pol-Pol_* as indicated). B) Lattice configurations upon addition of a polymer chain of length *L_polymer_* = 20 with *N_IC_* = 12 black monomers (black-black affinity: *w_IC-IC_* = —0.5) and *N_PC_* = 8 blue monomers (blue-red affinity: *w_PC-Pol_* = —0.5, adjustment see Supplementary Fig. 11) for the same range of *w_Pol-Pol_* values as in A). C) Lattice configurations for increasing sizes of the blue region (*N_PC_* as indicated, *w_Pol=Pol_* = —0.35 and *w_PC-Pol_* = —0.5). *N_IC_* as the difference of constant *L_polymer_* = 20 and varying *N_PC_*. All simulations have the same number of red particles, *N_Pol_* = 100, for adjustment of *N_Pol_*, see Supplementary Fig. 11A.

**Fig. 4:**
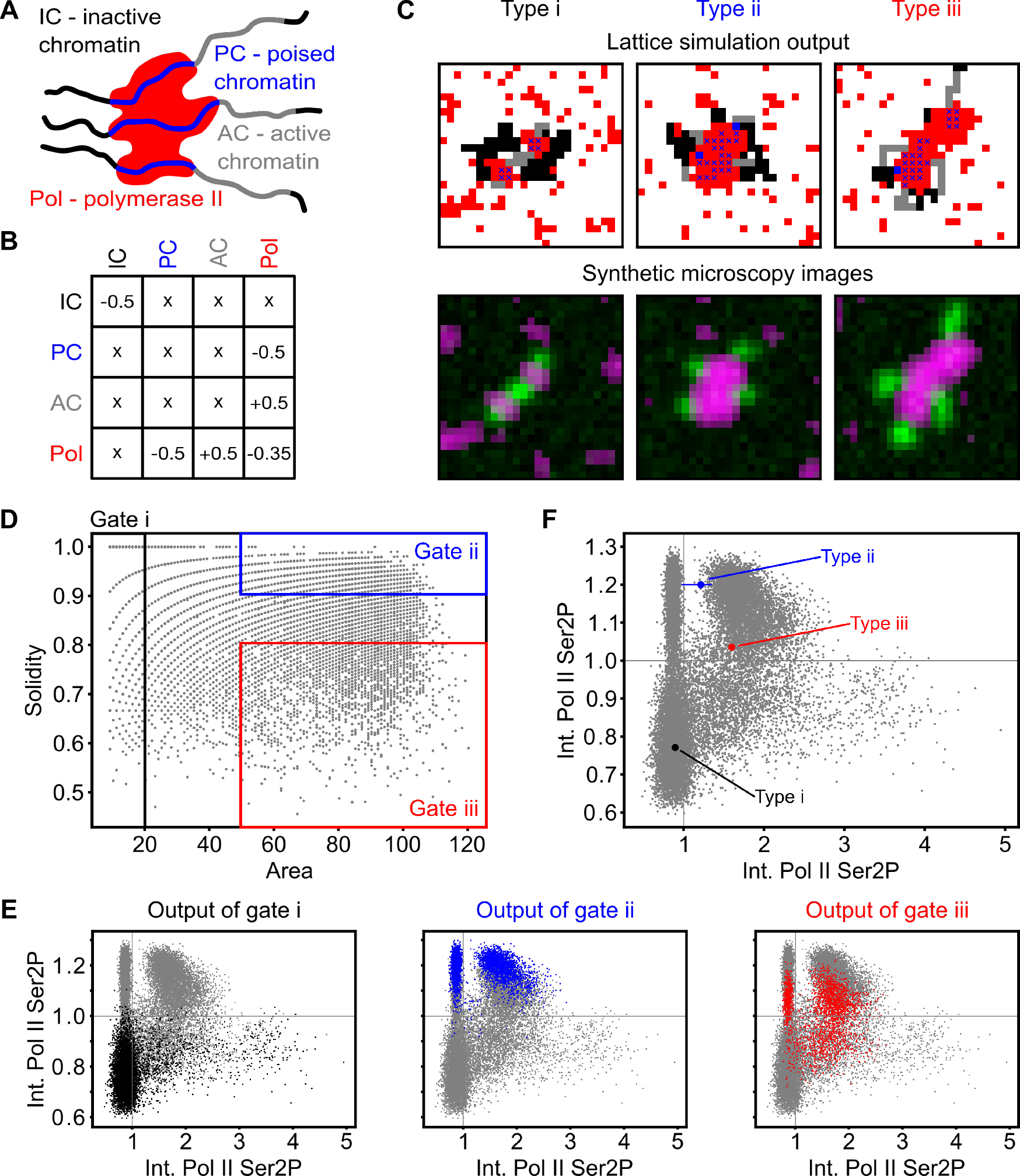
A lattice model with liquid phase wetting of promoter regions reproduces cluster morphologies. A) Sketch of cluster nucleation with all particle and polymer species involved in the model. B) Interaction matrix for different species in the lattice model. Affinity is represented by negative and repulsion by positive values. C) Example lattice configurations for all three morphological cluster types (i-iii), are shown as lattice simulation output and corresponding synthetic microscopy images. D) Area and solidity of individual clusters, with gate regions for the typical morphologies i-iii. Clusters were segmented based on Pol II Ser5P intensity (total 20401 clusters). The simulated configurations contained chains with different numbers of gray (*N_AC_* = 0,3,6), blue (*N_PC_* = 2,4,6,8) and black (*N_IC_* depending on *N_AC_* and *N_PC_*) monomers, with a total polymer length *L_polymer_* = 20. 6525 clusters in gate i, 4065 clusters in gate ii, 2326 clusters in gate iii. E) The Pol II Ser5P and Pol II Ser2P intensities (mean intensity across all pixels inside a given cluster’s segmentation mass) of the clusters in the gates i-iii are plotted in color over the entire ungated cluster population (light gray). F) The median of the Pol II Ser5P and Pol II Ser2P levels of the gated clusters in panel D is plotted over the ungated population of clusters. Each cluster type is plotted with 95% bootstrapped confidence intervals in Pol II Ser5P and Pol II Ser2P direction (10,000 resamples). For types i and iii, the confidence interval is hidden by the median data point. Mean Pol II Ser5P and Pol II Ser2P intensities were scaled by the median value for all time steps.

To introduce the effect of transcription elongation, we placed additional “gray” regions into the polymer chains directly adjacently to blue regions (referred to as AC, for active chromatin, see Fig. 4A). This placement reflects the location of gene bodies immediately adjacent to promoter regions. Based on the observation that elongating Pol II is excluded from droplets enriched in recruited Pol II^18–20^, we introduce repulsion between gray subregions and red particles (*w_AC-poi_* > 0, Fig. 4B). Finally, inactive chromatin was previously shown to segregate from other nuclear components, which was captured by self-affinity of these inactive regions (*w_IC-IC_* < 0)^25,33^. The inactive chromatin regions (“black” polymer stretches) were placed adjacently to the blue-gray domain. Simulations with different numbers of blue and gray monomers placed on the polymer chains produced configurations that resemble the cluster types i-iii seen in our microscopy images (Fig. 4C). To produce synthetic Pol II Ser5P and Pol II Ser2P intensity images, we applied a limited resolution filter and detector noise to the lattice distribution of red and gray particles, respectively (Fig. 4C). These synthetic microscopy images allowed us to apply the same morphology-based gating into type i, ii and iii clusters as in the real microscopy data (Fig. 4D). By this analysis, we found a similar relationship of Pol II Ser5P and Pol II Ser2P levels to cluster type as for our experimental data (Fig. 4E,F). This agreement indicates that the proposed lattice model with liquid phase wetting of promoter regions can explain how Pol II recruitment and elongation influence Pol II cluster morphology.

### Surface-wetting simulations predict inhibitor effects on cluster morphology

Simulations of the LKMC model in the regime of surface wetting imply that the levels of recruited and elongating Pol II directly determine the morphology of a given cluster. To test this implication, we applied two small molecule drugs, triptolide and flavopiridol, to inhibit Pol II recruitment and pause release in primary cell cultures of zebrafish embryos (Supplementary Figs. 12A and 14A). The effect on phosphorylation levels of Pol II clusters was in line with expectations^34^: triptolide reduced the Pol II Ser5P and Pol II Ser2P intensities, while flavopiridol reduced Pol II Ser2P levels (Fig. 5). Importantly, the inhibitors had visible effects on cluster morphology (Supplementary Fig. 13A).

**Fig. 5:**
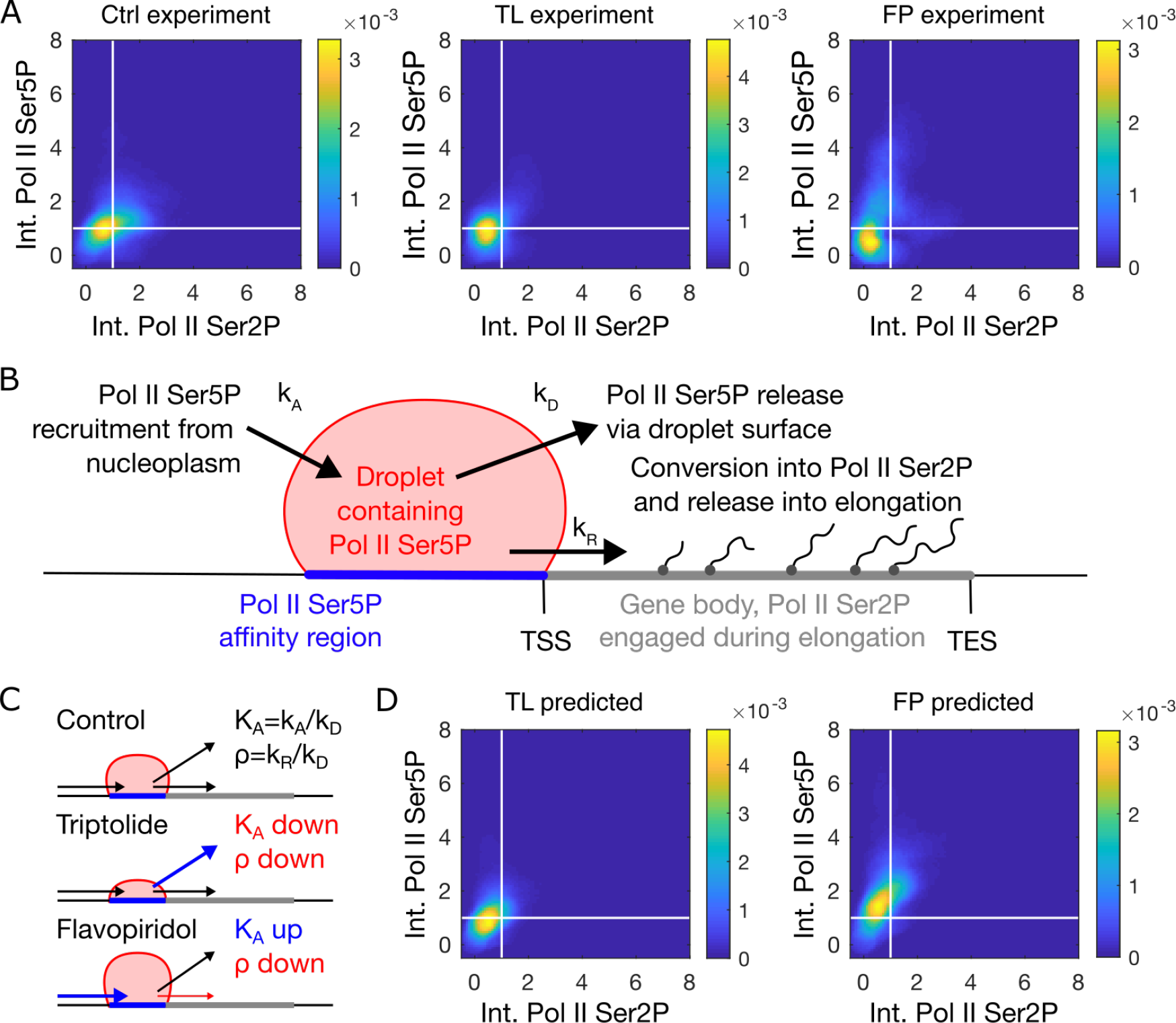
Characterization of kinase inhibitor effects on Pol II cluster formation using a Pol II compartment model. A) Mean Pol II Ser5P and Pol II Ser2P intensity distribution of Pol II clusters in data recorded from primary zebrafish cell cultures treated with DMSO control media (Ctrl), triptolide (TL), or flavopiridol (FP) for 30 minutes. B) Sketch of the mathematical model that was applied to reproduce changes in Pol II Ser5P and Pol II Ser2P intensities. A droplet containing Pol II Ser5P is the central model compartment, to which Pol II Ser5P is recruited (rate coefficient *k_A_*), and released either via the compartment boundary (rate coefficient *kp*) or by transition into transcription elongation (rate coefficient *k_R_*). C) Parameter changes used to reproduce the intensity distribution in the inhibitor-treated samples. D) Intensity distributions were reproduced based on the mathematical model and the control-treated intensity distribution. For every cluster in the control condition, the parameter values (*I,C*) were calculated, and subsequently transformed into altered Pol II Ser5P and Pol II Ser2P intensities by multiplication of *K_A_* and *ρ* with a prefactor representing the inhibitor treatment. After triptolide treatment, *K_A_* and *p* were 0.72 and 0.58 fold their control values, respectively. After flavopiridol treatment, *K_A_* and *p* were 1.18 and 0.76 fold their control values, respectively. The parameter estimation was based on minimizing the root-mean-square difference between predicted and measured Pol II Ser5P and Pol II Ser2P intensity distributions, see Supplementary Fig. 15.

To understand what changes need to be made in the parameters of the LKMC model to account for inhibitor treatment effects, we first rationalized the inhibitor-induced changes in Pol II Ser5P and Pol II Ser2P levels with a simple mathematical model. This mathematical model is distinct from the LKMC simulations and describes a liquid phase compartment with incoming fluxes (recruitment) and outgoing fluxes (release) of Pol II Ser5P in the form of rate equations (Fig. 5B, for details see Methods). By fitting this model to the observed changes in Pol II phosphorylation, we could capture the effects of triptolide and flavopiridol (Supplementary Fig. 15). Specifically, the effect of triptolide is best explained by a reduction in the Pol II Ser5P affinity to the droplet (*K_A_*) and a reduction in the ratio of pause release into elongation over release through the droplet surface (*p*) (Fig. 5C). The effect of flavopiridol is best explained by an increase in *K_A_* and a reduction in *ρ*. We used these modified *K_A_* and *p* values to transfer the Pol II Ser5P and Pol II Ser2P intensity distributions obtained from control-treated cells into predicted distributions from inhibitor-treated cells (Fig. 5D). These predicted distributions match the experimentally observed distributions, suggesting that the simple one-compartment model captures the changes in Pol II recruitment and release upon inhibitor treatment.

To predict inhibitor-induced changes of cluster morphology, we introduced parameter changes into the LKMC model that reflect the changes described by the compartment model. The simplest approach to mimic the reduction of *K_A_* and *ρ* detected for triptolide is by an increased release of Pol II from the liquid phase compartment that forms at blue regions. This was achieved by reducing Pol II self-affinity (*w_Pol-Pol_*) and surface affinity (*w_PC-Pol_*) (Fig. 6A). Simulations with these changes exhibited a lowered solidity and frequent fragmentation of clusters that led to a larger overall number of clusters (Fig. 6B). The increase of *K_A_* and reduction of *ρ* for flavopiridol was implemented as removal of all gray parts (*N_AC_*) of the polymer chains (Fig. 6A). This change reflected the marked reduction of elongating Pol II due to decreased *ρ*, and also resulted in an increase of cluster size representative of higher droplet affinity *K_A_* (Fig. 6A). Simulations with these changes exhibited an increased solidity, and an overall lower number of clusters (Fig. 6B). The predicted changes in solidity and overall cluster number were confirmed by our experimental data (Fig. 6C, for a full analysis of cluster properties, see Supplementary Figs. 14A and 12A). Considering that the model is based on affinity differences due to Pol II phosphorylation, it is expected that inhibitors that perturb transcription by other pathways should not be captured by the model. We therefore applied actinomycin D, which inhibits transcription by intercalating with DNA. While changes in phosphorylation did occur (decreased Pol II Ser2P, Supplementary Fig. 14A), the observed effect on morphology (i.e., decreased solidity, Supplementary Fig. 14A) is different from the model prediction (i.e., increased solidity). Taken together, these observations suggest that the model has strong predictive power for inhibitors that directly perturb phosphorylation of Pol II.

**Fig. 6:**
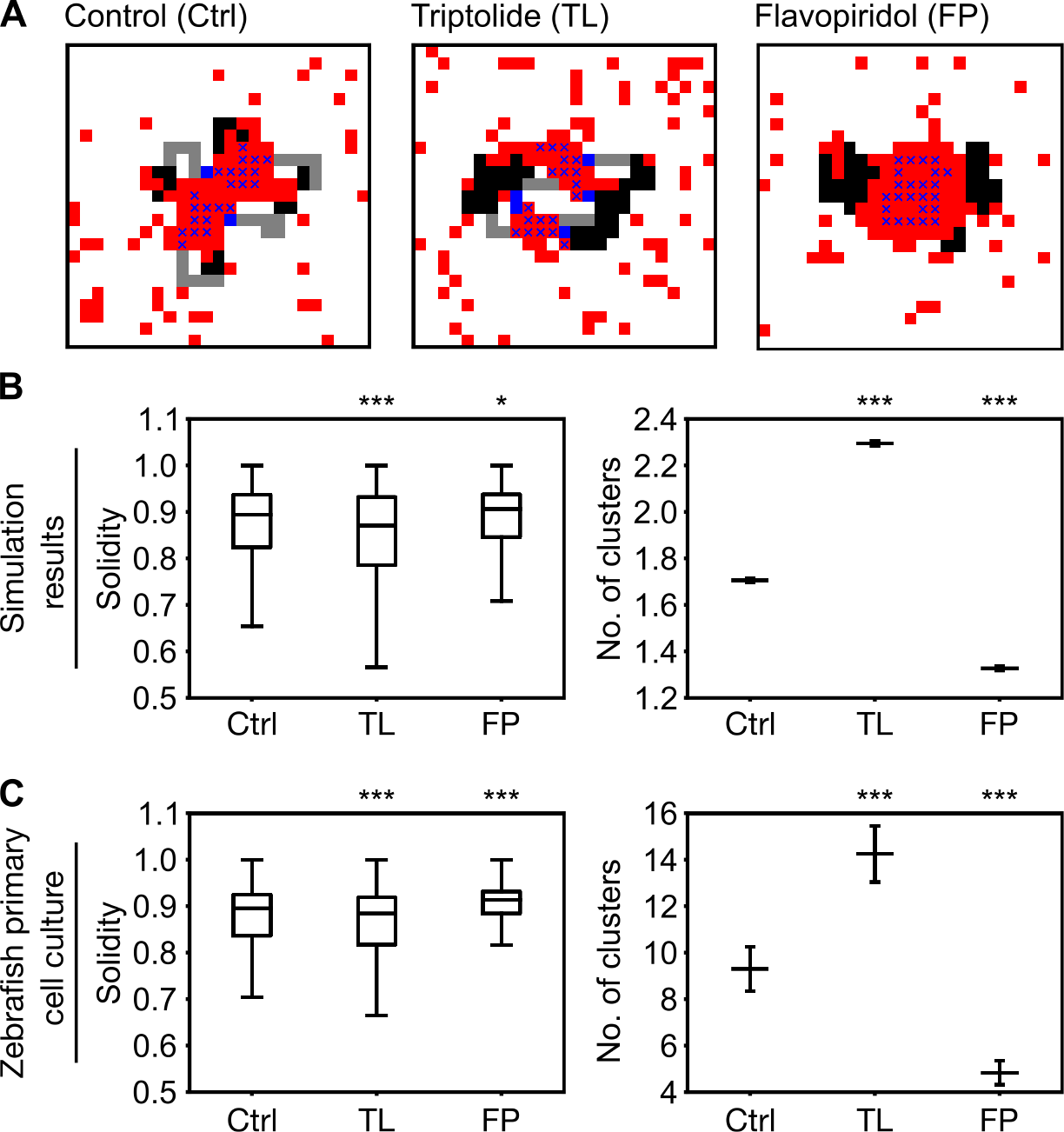
Liquid phase wetting simulations predict the effect of transcription inhibitors on cluster morphology. A) Examples of lattice configurations obtained from simulations with parameter changes that represent inhibitor treatment. For the control treatment (Ctrl), a range of *Npc* = 2,4,6,8 blue monomers per chain and *N_AC_* = 0,3,6 gray monomers per chain (total length *L_polymer_* = 20) was used; Pol II affinity parameters were assigned standard values (*w_Pol-Pol_* = —0.35, *w_PC-Pol_* = —0.5). For the triptolide treatment (TL), the same parameter range for *N_PC_* and *N_AC_* were used, and Pol II affinities were reduced (*w_pol-pol_* = —0.25, *w_PC-pol_* = —0.25). For the flavopiridol treatment (FP), no gray monomers were included (*N_AC_* = 0 throughout), Pol II affinities were unchanged. B) Cluster solidity and number of distinct clusters per simulation. Quantification was based on synthetic microscopy images, Ctrl contained 12,000 evaluated images, TL 12,000, and FP 4,000. Solidity is median with boxplots, number is mean±SEM; *** indicates *p* < 0.001, * indicates *p* < 0.05 (*p* < 0.0001, and *p* = 0.03 for solidity, *p* < 0.0001 and *p* < 0.0001 for number of clusters; permutation test for differences from control, resampling *n* matched to experiments in panel C, Bonferroni-corrected). C) Cluster solidity and number of distinct clusters per cell nucleus obtained from primary zebrafish cell cultures. *** indicates p<0.001 (*p* = 0.007, and *p* < 0.0001 with *n* = 1514,1631,703 clusters for solidity, *p* = 0.004 and *p* = 0.0002 with *n* = 402,294,150 nuclei for number of clusters; permutation test for differences from control, Bonferroni-corrected, data obtained from three independent sets of experiments). Full assessment of cluster and per-nucleus properties, see Supplementary Figs. 14A and 12A.

We proposed that cluster morphology is shaped by liquid phase wetting and dispersal. These processes should, in essence, apply independently of specific circumstances, for example, the specific cell type. We thus repeated the inhibitor experiments in a human cell line (THP-1, undifferentiated, Supplementary Fig. 13B). While clusters were smaller and fewer in number, the changes in phosphorylation levels and cluster morphology upon triptolide and flavopiridol treatment directly corresponded to those in zebrafish cells (Supplementary Fig. 14B). Again, the application of actinomycin D resulted in a morphological change (increased solidity) that could not be explained by changes in phosphorylation levels (no change, Supplementary Fig. 14B). These results demonstrate that our conclusions can be transferred to at least one other cell type.

## Discussion

In this study, we investigated how the morphology of clusters enriched in recruited Pol II is determined by the levels of recruited and elongating Pol II. Our findings indicate that these clusters behave like droplets formed by wetting of chromatin by a liquid phase enriched for recruited Pol II. Chromatin regions containing elongating Pol II are excluded from this liquid phase, resulting in a dispersed droplet morphology. Previous work has described how genomic regions can influence the coarsening of droplets that form by canonical LLPS, meaning that droplets would spontaneously form also in absence of these regions^35,36^. In other cases, formation of clusters requires transcriptional activity at specific genomic loci, suggesting a more complex scenario than canonical LLPS^10,13,14^. Recent *in vitro* studies of condensate formation on DNA suggest that liquid phase condensation that requires a condensation surface can be accurately described as wetting by a sub-saturated liquid phase^23,24^. Our work in an embryonic model system suggests that the formation of clusters enriched in recruited Pol II might proceed similarly by liquid phase wetting with chromatin as a condensation surface.

Previous studies of Pol II clusters have already revealed dispersed morphologies of Pol II clusters, which markedly deviate from the droplets with smooth surfaces expected for canonical LLPS ^13,22^. Complex morphologies of phase-separated droplets can, for example, occur in multi-component condensates^37^. Such a multi-component perspective might also be applied to Pol II clusters, considering that the phosphorylation states of recruited and elongating Pol II were found to spatially exclude each other ^15,18–20^. Our theoretical model explains how dispersal of droplets could result from exclusion of chromatin harboring elongating Pol II from the liquid phase enriched in recruited Pol II. Recruitment and elongation would thus occur in close-by, though spatially segregated compartments, which are tied to each other through common chromatin templates. This idea has been proposed for Mediator-Pol II clusters in mouse embryonic stem cells (mESCs)^13^, and our work might provide a theoretical model for such a compartmentalization. It is also in line with the recent observation of chromatin domains that move as connected units^38–40^.

Our model of cluster formation is coarse-grained in nature, compressing the molecular reality of the biological cell into a small number of generalized components. For example, the control of recruitment, initiation, pausing, and pause release proceeds along numerous steps, and the list of involved regulatory factors is continuously expanding^20,41–43^. Two key regulators that were identified in zebrafish embryos are p300 and BRD4, which are associated with the histone 3 lysine 27 acetylation (H3K27ac) active chromatin mark^26,44^. H3K27ac has also been identified as a crucial requirement for pause release into elongation^45^. Additional assessment of these regulators and histone marks with respect to the morphology of Pol II-enriched clusters should thus provide a more comprehensive understanding.

The immunodetection approaches used in this study could be further expanded by sequence-specific staining of genomic regions or RNA transcripts. Such additional staining could connect the morphology of Pol II clusters to the 3D organization of chromatin within and in the neighborhood of these clusters^46,47^. Such an assessment would touch upon the long-standing question whether genes get recruited to Pol II clusters and subsequently activate, or if instead transcription induction establishes clusters instead^8,48,49^. In the case of one gene in mESCs, *Esrrb,* Pol II clusters were only transiently visited^13^. In contrast, in zebrafish embryos, activation of the microRNA miR-430 cluster drives the *de novo* establishment of Pol II clusters following cell division ^25,44,50^.

In our time-lapse recordings during interphase, Pol II clusters maintained different types of morphologies over ten minutes and longer. It is in line with previous observations that nuclear bodies and their three-dimensional organization, once established after cell division, remain long-term stable^51^. A more fine-grained analysis of fluctuations of cluster morphologies might allow an assessment of how far active catalytic processes might result in characteristic non-equilibrium fluctuation signatures^52^. Also, transcription largely shuts down during mitosis in zebrafish blastula cells during cell division^25^. It would therefore be interesting to assess how Pol II clusters are reestablished following cell division. The reshaping and reestablishment of Pol II foci have recently also been connected to ATP-dependent catalytic processes associated with nuclear actin and myosin^53,54^. Taken together, there is indication of a number of catalytic and mechanochemical processes that might contribute to the establishment and changes of Pol II cluster morphology and await further investigation.

We categorized cluster morphologies based on size (area) and one characteristic that captures the apparent degree of dispersal (solidity). This characterization was sufficient to establish a direct relationship between Pol II phosphorylation and cluster morphology. However, the observed morphologies warrant a more comprehensive morphological characterization. Especially those clusters with complex morphologies – resembling for example donuts, croissants, or jalebis – provide an intriguing variety of shapes to support further work. Amongst these, the donut-shaped clusters that persist for ten minutes and longer (Supplementary Fig. 7) are most intriguing, and might be connected to the looping of transcription termination sites to the promoter region^55,56^.

According to our model, Pol II recruitment relates to the establishment and elongating Pol II to the dispersal of macromolecular clusters. These findings correspond well with another recently proposed model, where RNA produced at gene regulatory elements supports the formation of condensates, whereas RNA produced during elongation of gene bodies can drive their dissolution^57^. Such an RNA-mediated role of elongation in dispersal of macromolecular assemblies is also in line with previous work implicating RNA in the unfolding of chromatin regions harboring transcribed genes^25,58,59^. Our particular focus on the morphology of macromolecular clusters closely assesses the interplay of these two opposing forces: cluster establishment by liquid phase wetting of regulatory chromatin regions, and the dispersal of clusters as a consequence of Pol II transitioning into elongation.

## Materials and Methods

### Zebrafish husbandry

Fish were raised and bred according to local regulations in the fish facility of the Institute of Biological and Chemical Systems. Embryos were produced by spontaneous mating. Embryos were dechorionated with Pronase, washed with E3 embryo medium, and subsequently kept in agarose-coated dishes in 0.3x Danieau’s solution at 28.5°C.

### Imaging of Pol II phosphorylation states in live zebrafish embryos

Covalently labelled Fab antibody fragments were injected into the yolk of dechorionated embryos at the single cell stage. Per embryo, 1 nl of Fab mix (0.2 *μ*l 1% Phenol Red, 1.5 *μ*l A488-labeled anti-Pol II Ser2P Fab, 2.3 *μ*l Cy3-labeled anti-Pol II Ser5P Fab, Fab stock concentration ≈ 1mg/ml) was injected. Embryos were mounted at the High stage in 0.7% low melting point agarose in 0.3x Danieau’s solution in ibidi 35 mm imaging dishes (#1.5 selected glass cover slips).

### Primary cell culture from zebrafish embryos

Fish embryos were collected in oblong stage and moved to low-retention microcentrifuge tubes. The embryos were deyolked through vortexing in deyolking buffer (55 mM NaCl, 1.75 mMKCl, 1.25 mM NaHCO3). Afterwards, 1 ml PBS (Dulbecco’s formulation) with 0.8 mM CaCl_2_ was added to the samples and incubated for 30 minutes. Inhibitors were introduced to PBS before distribution to individual culturing tubes. Samples were fixed by addition of 330 *μ*l of 8% Formaldehyde in PBS with 0.8 mM CaCl_2_ to each tube. Tubes were immediately spun down at 800 × g and left for 15 minutes at room temperature, then the liquid was replaced by 8% formaldehyde in PBS+CaCl_2_, left at room temperature for further fixation for at least 20 minutes.

### THP-1 cell culture

Undifferentiated cells from the human monocytic cell line THP-1 were generously provided by the Weiss laboratory, Institute of Biological and Chemical Systems, Karlsruhe Institute of Technology^60^. Cells were transferred into low-retention microcentrifuge tubes directly before experimental treatment, inhibitors were applied by spike-in and incubated for 30 minutes at room temperature, and fixation was carried out identically to primary zebrafish cell cultures.

### Inhibitor treatment

All inhibitors were resuspended in DMSO to recommended effective concentrations^34^. Flavopiridol hydrochloride hydrate (F3055, Sigma-Aldrich) was resuspended to a stock concentration of 12.5 mM, and diluted 1:12500 to an effective concentration of 1 *μ*M. Actinomycin D (A1510, Sigma-Aldrich) was resuspended to an initial concentration of 1 mg/ml, and diluted 1:200 to an effective concentration of 5 *μ*g/ml. Triptolide (T3652, Sigma-Aldrich) was resuspended to a stock concentration of 10 mM, and diluted 1:20000 to an effective concentration of 500 nM. The effectiveness of all inhibitors was verified on the basis of Pol II phosphorylation changes at the whole nucleus level (Supplementary Fig. 12).

### Immunofluorescence

Fixed cell cultures were processed for the entire immunofluorescence procedure in the low-retention microcentrifuge tubes in which they were cultured. Cells were permeabilized with 0.5% Triton X-100 in PBS for 10 min, washed three times with PBS with 0.1% Tween-20 (PBST), and blocked with 1 ml of 4% BSA in PBST for 30 minutes. Primary antibodies against Pol II Ser2P (abcam ab193468, rabbit IgG monoclonal, dilution 1:1000), Pol II Ser5P (clone 4H8, abcam ab5408, mouse IgG monoclonal, dilution 1:1000) and H3Ser28P (abcam ab10543, rat IgG monoclonal, dilution 1:10,000) were applied overnight at 4°C in 4% BSA in PBST. Secondary antibodies (Invitrogen A11037, Alexa Fluor 594-conjugated goat anti-rabbit IgG, dilution 1:1000; Invitrogen A11001, Alexa Fluor 488-conjugated goat anti-mouse IgG, dilution 1:1000; Invitrogen A21247, Alexa Fluor 647-conjugated goat anti-rat IgG, dilution 1:1000) were applied overnight in 4% BSA in PBST. Primary and secondary antibodies were removed by washing three times with PBST. After washing out the secondary antibodies, the samples were again fixed with 8% formaldehyde in PBS for 15 min for long-term retention of antibody staining. Samples were washed another three times with PBST, then mounted using 30 *μ*\ of Vectashield H-1000 supplemented with a 1:2500 dilution of Hoechst 33342 (stock concentration 20 mM) using selected #1.5 cover slips.

Animal caps of fixed whole embryos were obtained by fixing sphere stage embryos overnight at 4°C (2% formaldehyde, 0.2% Tween-20 in 0.3x Danieau’s embryo media). Animal caps were permeabilized in 0.5% Triton X-100 in PBS for 15 minutes room temperature, washed three times with PBST for 10 minutes, and blocked in 4% BSA in PBST for at least 30 minutes at room temperature. Primary antibodies against Pol II Ser5P (clone 4H8, abcam ab5408, mouse monoclonal IgG diluted 1:300) and Pol II Ser2P (abcam ab193468, rabbit monoclonal IgG, diluted 1:2500) were applied overnight at 4°C in 4% BSA in PBST. Secondary antibodies (Invitrogen A21206), Alexa Fluor 488-conjugated donkey anti-rabbit IgG, diluted 1:2000; Abberior 2-0002-011-2, goat anti-mouse IgG, conjugated with STAR RED, diluted 1:300) were applied overnight at 4°C in 4% BSA in PBST. Primary and secondary antibodies were removed by washing three times with PBST. After washing out the secondary antibodies, the samples were again fixed with 4% formaldehyde for 15 min for long-term retention of antibody staining. In most cases, these post-fixed embryo samples were free of yolk, and any remaining pieces of yolk were manually removed with fine forceps while transferring samples through three washes of PBST in glass dishes. The deyolked animal caps were mounted in TDE-O (Abberior) using selected #1.5 cover slips.

### Instantaneous Structured Illumination Microscopy (instant-SIM)

Microscopy data from live whole embryos and inhibitor-treated, fixed cells were recorded using a VisiTech iSIM high-speed super-resolution confocal microscope based on the instant-SIM principle^27^. The microscope was built on a Nikon Ti2-E stand. A Nikon Silicone Immersion Objective (NA 1.35, CFI SR HP Plan Apochromat Lambda S 100XC Sil) was used for live imaging, a Nikon Oil Immersion Objective (NA 1.49, CFI SR HP Apo TIRF 100XAC Oil) was used for fixed cell imaging. Excitation lasers at 405, 488, 561 and 640 nm were used, illumination and acquisition settings were kept constant across all samples of a given experimental repeat. Color channels were recorded on two cameras simultaneously for increased speed during live imaging, and sequentially to avoid cross-talk during fixed cell imaging.

### STimulated Emission Double Depletion (STEDD) microscopy

Microscopy data from animal caps of fixed whole embryos were recorded using a custom-built STEDD microscope, as previously described^30^. The STEDD principle allows suppression of low frequency image components as well as out-of-focus light and aberrant signal from reexcitation effects^28,29^. Here, STEDD-resolved images were recorded using excitation by a 640 nm pulsed laser (675 nm detection band-pass filter), depletion by a titanium-sapphire depletion laser tuned to 779 nm, focused though an oil-immersion objective (HCX PL APO CS 100 ×/1.46, Leica). The confocal image was acquired in an additional scan in the same focal plane, using a 473 nm excitation laser (520 nm detection band-pass filter) without additional depletion.

### Image analysis – instant-SIM microscopy data

Single time point z stacks were maximum-projected in FIJI, including the 25 slices with best image quality in a given stack^61^. The further 2D analysis was implemented as a Cell Profiler pipeline^62^. Specifically, a two-step approach was used, where first cell nuclei and subsequently Pol II clusters inside nuclei were segmented based on the Pol II Ser5P Fab signal. Nuclei segmentation masks were obtained by global Otsu thresholding. Cytoplasmic masks were generated by outward dilation (25 pixels) from the nuclear masks. Pol II clusters inside nuclei were obtained by enhancing the Pol II Ser5P channel (speckle enhancement) and global robust background thresholding (5.5 standard deviations). For each cluster, the mean Pol II Ser5P and Pol II Ser2P intensities (cytoplasmic background subtracted on per-nucleus basis), cluster area, and cluster solidity were extracted. The mean intensity of Pol II Ser5P and Pol II Ser2P in the nuclei, cytoplasm and in single clusters were measured. The geometric properties – solidity, area – gwere measured for each cluster. All clusters smaller than 4 pixels were discarded. Further data processing and graph preparation was done in Python^63^. Data from fixed cells was analyzed in the same way, but as a first step used the additional color channels with DNA and H3Ser28P signal to establish Otsu-threshold masks for nuclear segmentation and prophase exclusion, respectively. In the robust background segmentation of Pol II Ser5P clusters, 6.5 standard deviations were chosen for zebrafish primary cell cultures, 8 standard deviations for THP-1 cell cultures. A comparison of our analysis based on twodimensional, maximum-projected images with an analysis of full three-dimensional stacks showed a good correlation between both approaches (Supplementary Fig. 8).

### Image analysis – STEDD microscopy data

Nuclear segmentation masks were obtained by Gaussian blur (*σ* = 1.2 *μ*m) and Otsu thresholding. Pol II Ser5P clusters and Pol II Ser2P spots were segmented by local background subtraction (Gaussian blur image with *σ* = 0.4 *μ*m subtracted), followed by global robust background thresholding (0.5 and 0.25 Standard Deviations). For each Pol II Ser5P cluster, mean intensity, area, and solidity were extracted. For Pol II Ser2P spots, only the mean intensity was extracted due to the lower confocal resolution relative to the STEDD data. For the relative placement analysis (Fig. 2F), 8-neighborhood distances were used. The analysis was carried out using MatLab.

### Statistics

Box plots conform to standard practice (median, quartiles as boxes, range as whiskers, outliers removed outside of 1.5 times interquartile range extension). Statistical significance was indicated for differences of mean relative to the control condition, permutation test with Bonferroni correction; *, **, and *** indicate p<0.05, p<0.01, and p<0.001, respectively.

### Lattice model of cluster formation

Monomeric particles that represent Pol II Ser5P and polymeric chains that represent chromatin were placed on a 25 × 25 two-dimensional grid. Lattice configurations were obtained using a rejection-free Lattice Kinetic Monte Carlo (LKMC) algorithm: single particles can move to all eight nearest neighbours, while polymer chains can only move according to the Verdier-Stockmayer move-set (end-bond flip, kink-jump, and crankshaft move), which ensures chain connectivity^32,64,65^. Synthetic microscopy images were obtained by converting lattice distributions of Pol II monomers and lattice distributions into matrices with values 0 (unoccupied) or 1 (occupied), which were blurred with a Gaussian kernel (*σ* = 1). For both synthetic color channels, artificial detector noise was added in the form of Poisson-distributed random numbers (mean *λ* = 5, divided by 100 before addition). Objects corresponding to Pol II Ser5P clusters were obtained from these blurred distributions using a segmentation threshold 0.35 followed by connected-component analysis (2-connectivity, components with less than 10 pixels were excluded). Distributions of area, solidity, and intensities of these objects were obtained by sampling simulations at different time points (ergodic sampling). The numerical simulation and the analysis of synthetic images were carried out with Python. For details, see Supplementary Materials.

### Single compartment model for Pol II recruitment and release

The central part of the model is a compartment of volume *I*, which represents a Pol II Ser5P droplet formed at the promoter regions. The dynamics of the volume of this compartment is described as

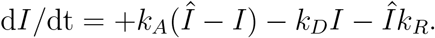

This compartment has a finite capacity *Î* to recruit Pol II Ser5P, and this recruitment is assumed to happen in a single step with rate coefficient *k_A_*. Pol II Ser5P can be released back to the cytoplasm through the droplet surface. Again, we assume a single step reaction with rate coefficient *k_D_*. Pol II Ser5P can also undergo initiation to the promoter, and subsequent conversion into Pol II Ser2P and release into transcript elongation. Here, we take the Pol II Ser5P concentration inside the droplet as approximately constant, independent of droplet volume *I*, so that this reaction is modeled with a constant rate coefficient *k_R_*^1,10,66^. However, the global rate of this reaction is proportional to the number of promoter sites that can transform recruited into elongating Pol II, so this term is proportional to *Î*. Considering that the clusters exhibit long-term stable morphologies, we calculate *I* at steady state (d*I*/dt = 0):

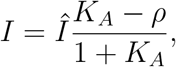

where *K_A_* = *k_A_/k_D_* is the effective affinity of Pol II Ser5P for the compartment (in the *k_R_* = 0 case), and *ρ* = *k_R_/k_D_* the ratio of pause release vs release across thee droplet interface. The elongating polymerase persists in the gene body for an average time *T_elong_*, resulting in a steady state level of Pol II Ser5P of

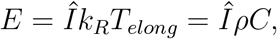

with *C* = *k_D_T_elong_*. The parameters *k_A_* ≈ 1/(54 s), *k_D_* ≈ 1/(7s), *k_R_* ≈ 1/(150 s) are known^45^. The length of genes at the sphere stage of zebrafish embryo development is ≈ 10 kb^67^, and the elongation rate ≈ 2kb/min^68^, resulting in *T_elong_* ≈ 5 min. We thus set *K_A_* = 0.1 and *C* =1. *I* and *K_R_* can be calculated for single clusters using the measured Pol II Ser5P and Pol II Ser2P intensities as the values for *E* and *I*, respectively. The distribution of *ρ* resulting from this approach is centered on the predicted value of 0.05 (Supplementary Fig. 15). *Î* as an effective parameter cannot be compared to any experimental measurement we are aware of.

## Acknowledgments

We thank Haruka Oda, Yuko Sato, and Hiroshi Kimura for Fabs, Susanne Fritsch-Decker and Carsten Weiss for THP-1 cells, Martin Bastmeyer and Carmelo Ferrai for discussion. The work was supported by the Helmholtz program Biointerfaces in Technology and Medicine (BIFTM). G.U.N. acknowledges financial support by the Helmholtz program Science and Technology of Nanosystems (STN), Max Planck School of Photonics (MPSP), and the Karlsruhe School of Optics and Photonics (KSOP). T.K., I.M., V.Z., and L.H. acknowledge financial support by the German Research Foundation (Priority Program 2191, Molecular mechanisms of functional phase separation, Project 419138152). The authors declare that they have no conflict of interest.

## Supplementary Materials

### Lattice Model

#### Lattice kinetic Monte Carlo model

We describe the Pol II cluster morphologies observed in zebrafish experiments using a simple physical model, which is limited to the most essential components: RNA polymerase II (Pol II, monomeric particles) and chromatin (linear polymeric chain) with different subregions. To obtain spatial configurations of this system, we used a rejection-free Lattice Kinetic Monte Carlo (LKMC) algorithm. LKMC algorithms, generally speaking, are suited to simulate coarse-grained stochastic non-equilibrium systems. The rejection-free algorithm (Fig. 10A) is similar to the Gillespie algorithm^69^. At the beginning of a simulation, by checking the system configuration and nearest neighbours of every particle within the system, a rate catalog with all possible transitions is created, providing also the total system rate as the sum of all transition rates. This initial cataloging step is followed by a Monte Carlo (MC) routine that is repeated *N* times. During each step, one of the previously defined transitions is randomly selected while associating transitions with a higher rate with a higher likelihood of occurrence. The transition is then performed and changes the system state. This is followed by a local update of the possible transitions in the affected lattice area, the total rate of the system, and the system time. This simulation paradigm has been used to model surface catalysis processes^70^ and slip-link DNA systems with DNA polymers and ring proteins^32^. The initialization of the simulations proceeds similar to the latter work, but instead of ring proteins uses the single Pol II complexes in addition to the polymers.

#### Initial configuration

Chromatin is modeled as a connected polymer chain with different internal states: inactive (black), poised (blue), and active (gray). Exclusion from occupied volume is assumed, so that chains can only undergo a limited type of moves that maintain chain connectivity. The Pol II Ser5P particles (single lattice sites, red) can freely diffuse in space and interact with different affinities *W_i_* with the DNA polymer and other Pol II Ser5P particles. The different interspecies affinities of Pol II to specific subregions of DNA allow us to study the formation of Pol II clusters in the framework of microphase separation. At the beginning of a given simulation, the Pol II particles are randomly distributed on the lattice. The DNA polymer was placed in different initial configurations (e.g. single chain, cross of four chains, four parallel chains, four chains organized as random walks). The monomers making up chromatin chains were assigned to the different colors, giving contiguous sections of black polymer (number of monomers: *N_IC_*), blue monomers (*N_PC_*) and gray monomers (*N_AC_*). The number of red particles (*N_Pol_*) can be varied.

#### Pol II and DNA move set

After initialization and each time step, the rate catalog is updated. To find all possible transitions of the system for the rate catalog, we first have to define the allowed move set for every species. The Pol II Ser5P particles are allowed to move to one of its eight nearest neighbors in one MC step (Figure 10D, left). The polymer is simulated as a connected and self-avoiding chain. We therefore use the common Verdier-Stockmeyer move set, consisting of end-bond flip, kink-jump, and a crankshaft move (Fig. 10D, right)^32,64,65^. Movements to positions outside the lattice are not considered.

The end-bond flip can occur only for the first or last monomer of the polymer, and moves the first/last monomer to any lattice site neighboring the second/second-last particle of the polymer, changing the angle between the old and new positions by 90 degrees. For a kink jump, a monomer is moved to the opposite side of a corner formed by the preceding monomer, the monomer itself, and the subsequent monomer. To reach ergodicity of the system, a third move, called the crankshaft move, is added^64^. While the end-bond flip and the kink jump move only one monomer, the crankshaft move changes the position of two successive monomers.

#### Rate catalog

To draw up the rate catalog, we first browse all Pol II Ser5P positions within the lattice and test the occupation of its eight nearest neighbors. Those which are not occupied by the same species are possible directions for transitions within the system. Together with the defined rate coefficient, *k_p_* = 0.1, and the Arrhenius equation,

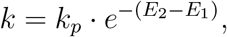

the rate for each transition can be determined. The energies before (*E*_1_) and after (*E*_2_) are stated in units of *k_B_T*, and are determined under consideration of the different interspecies affinities between the swapped particles and their nearest neighbors (Fig. 10C). Where a nearest neighbor position is located outside the lattice, no energy contribution is added for that neighbor. For every Pol II Ser5P transition, the old and new position and also the rate for the transition are added to the rate catalog. The transition rate is also added to the overall system rate *k_total_*.

As a second step, the same procedure is carried out for moves of the polymer chain that represents chromatin. Initially, the position within the polymer and also the position of the previous and subsequent monomers are checked. Depending on the position and configuration, only certain movements of the monomer are possible. As with the polymerases, it must then be checked whether the possible new position resulting from the move is already occupied by another monomer. If this is not the case, the Arrhenius equation and the rate coefficient for a single move (same as *k_kink_*) are used to determine the rate of the transition. Since the crankshaft move affects two monomers at the same time, this coefficient is smaller compared to the other moves, 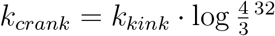. The polymer transitions are also added to the rate catalog. In addition to the positions and rates, the specific type of movement is added. Furthermore, for the crankshaft move both positions before and after the respective move are saved. Now all required properties of the system are defined and the main part of the Monte Carlo simulation can start.

#### Perform Move (MC-Step)

In this section, the core iteration step of the Lattice Kinetic Monte Carlo algorithm is described. At first, we draw a transition out of the previously defined rate catalog. This is done using a uniformly distributed random number *r*_1_ from the interval (0,1], the total rate *k_total_*, and the tower-sampling method (Fig. 10E). The transition rate *k*\* is calculated by *k*\* = *r*_1_ · *k_total_*. By looping through all transitions within the catalog and summing up each individual rate *k_i_* we identify a transition *j* with rate *k_j_* such that the relation

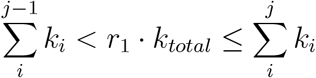

is fulfilled. The chosen transition is then performed and the system lattice is updated. The system time is also updated by adding

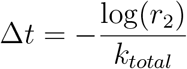

with a second random number *r*_2_ from (0,1] to the current system time *t*^32^. This routine is performed *N* times.

#### Local Update

At the end of each of the above iteration steps, the system rate catalog has to be updated. In the simplest approach, we simply delete the previous rate catalog and completely recalculate it. While formally correct, more computationally efficient alternative approaches are available^32^. Position changes in the simulation are only made in a confined area surrounding the particle that is moved. Accordingly, only the transitions falling within this area need to be updated. We therefore determine all transitions leading to a position within this area, or whose origin is within this area and update only these transitions. Since the longest transition within the system is performed by the crankshaft move with a distance of two lattice sites, the local update has to be performed in the region [*x* ± 4,*y* ± 4] (Fig. 10F).

#### Parameter Adjustment

Besides Pol II self-affinity *w_Pol-Pol_* (Fig. 3), we also adjusted the amount of Pol II Ser5P particles, *N_Pol_,* and Pol II Ser5P affinity to poised chromatin, *w_Pc-Pol_*. To adjust *N_Pol_*, we add a single polymer chain of length *L_Polymer_* = 20 with black and blue subregions to the simulation (Supplementary Fig. 11A). We use the previously determined affinity *w_Pol-Pol_* = −0.35, and assign *w_PC-Pol_* = −0.5 as a preliminary value. We then varied the amount of Pol II particles (*N_Pol_* = 10,25,50,100,200). Since we planned to perform simulations with four or more polymers and different lengths of the blue regions, we choose *N_Pol_* = 100 so as to provide enough material for cluster formation (Supplementary Fig. 11A). To assess the preliminary *w_PC-Pol_* = −0.5 value, we performed simulations containing a single polymer chain (*L_Polymer_* = 8) that consists only of blue subregions, using again *w_Pol-Pol_* = −0.35 and the previously determined amount of Pol II. As we vary *w_PC-Pol_* = −0.1, – 0.3, – 0.5, – 0.7, −1.0, we find that the only parameter for which not the whole polymer is covered by Pol II (blue still visible) is *w_PC-Pol_* = −0.1 (Supplementary Fig. 11B). For all tested values, cluster formation occurs, so that we continued using the preliminary value *w_Pol-Pol_* = −0.5, which falls in the middle region of the tested interval.

## Supplementary Figures

**Fig. 7:**
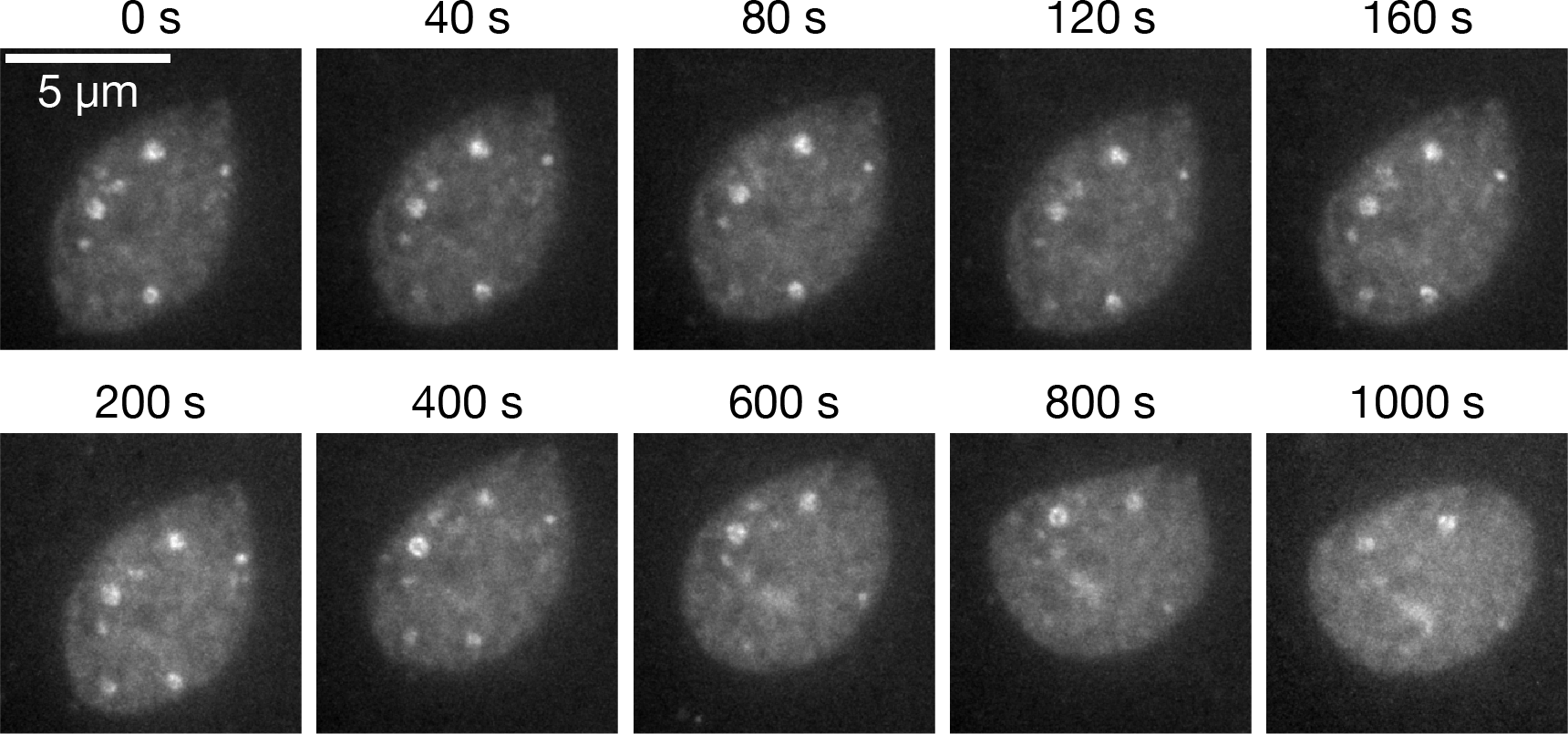
Time-lapse imaging of paused Polymerase II clusters. Clusters were visualized in live zebrafish embryos (sphere stage) using antibody fragments (Fab, Cy3-labeled) against the RNA polymerase II carboxyterminal domain serine 5 phosphorylation. The time-lapse images are maximum projections from a data set with 40 s time resolution signal. Images were chosen at selected time points to show short-term and long-term behavior. The same persistence of cluster morphologies was observed in time lapse sequences recorded from three embryos in parallel in the same experiment, and in a second independent repeat of this experiment.

**Fig. 8:**
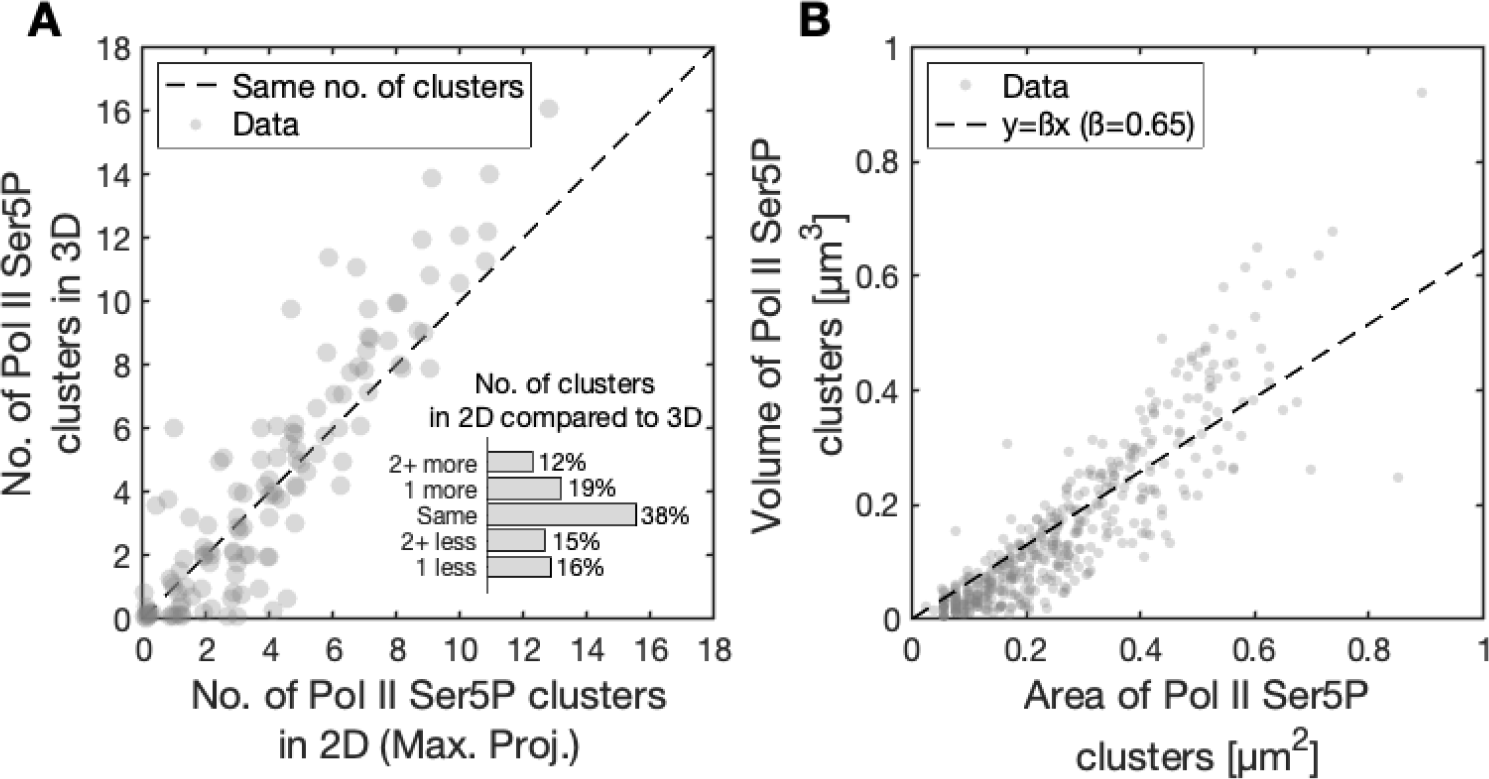
Comparison of 3D vs. 2D segmentation and analysis of polymerase clusters. **A)** To test whether our analysis of maximum-projected, two-dimensional (2D) microscopy data is representative of three-dimensional (3D) organization, we compare the number of Pol II Ser5P clusters detected per nucleus in both cases. A plot of the numbers from 2D segmentation based on maximum intensity projection (x-axis) and from 3D segmentation (y-axis) shows only small deviations from the dashed diagonal line (indicating perfect agreement). In the inset, we show the fraction of nuclei in which the number of clusters in the 2D and 3D methods are the same, differ by just 1 cluster, or by 2 and more clusters. The two methods result in a similar number of clusters: 38% of the nuclei have same number, and a total of 72% of nuclei differ in the 2D and 3D clusters at most by 1 cluster. The clusters were segmented using a global Robust Background Threshold (3 standard deviations for 2D projection and 4.5 standard deviations for the 3D image) on the masked images. **B)** We compare the area and volume of the clusters obtained from the 2D segmentation and 3D segmentation, respectively. For each nucleus, we obtained a set of common clusters in 2D and 3D by comparing their x, y positions (ignoring the z position of the 3D clusters) and picking those that overlap. For these clusters, a direct comparison is possible, and we can see that the volume and area are highly correlated. A linear fit results in a slope of 0.65, which gives an effective mean radius of about 0.5 *μ*m if the clusters were spherical. Overall, this comparison of 3D and 2D Pol II Ser5P clusters shows that 2D segmentation using maximum intensity projections results in a similar number and size of clusters when compared to a 3D analysis, justifying our approach of using the 2D method for our main analysis.

**Fig. 9:**
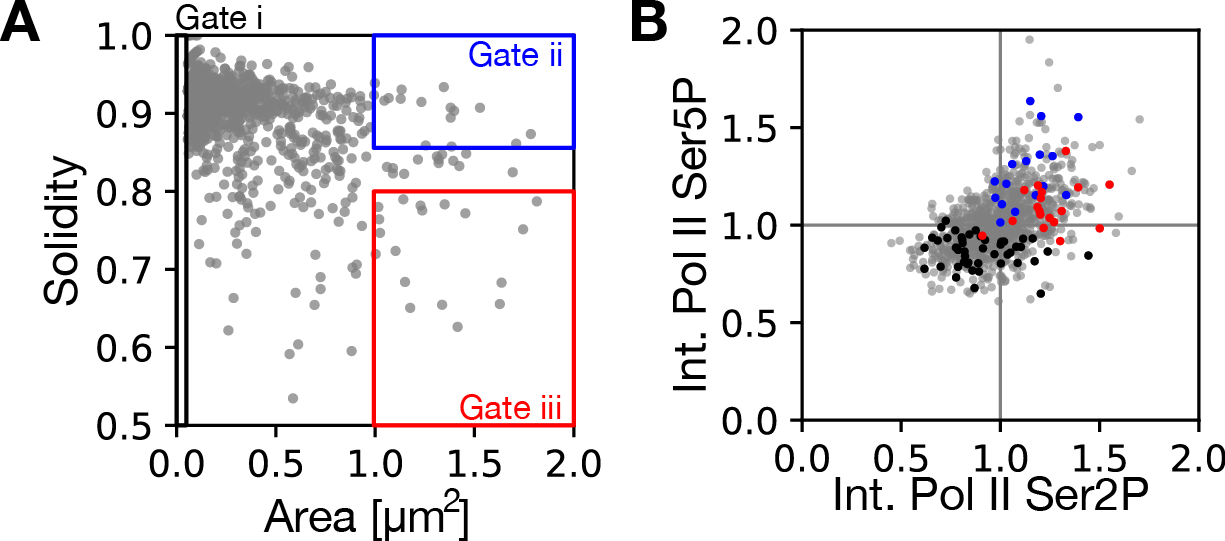
Quantification of cluster morphologies and phosphorylation levels from live embryos. Cluster properties were extracted from a zebrafish embryo (sphere stage) during live imaging using antibody fragments (for details, see Materials and Methods). A) Area and solidity of the individual clusters, with gates showing the type i, ii and iii clusters. B) Phosphorylation levels (serine 2 and serine 5 phosphorylation) of the individual clusters were monitored by mean fluorescence intensity. Clusters belonging to one gate tend to have similar Pol II Ser2P and Pol II Ser5P intensities. The values are normalized to the median value. Number of nuclei: *n* = 109. Number of clusters: *n* = 959.

**Fig. 10:**
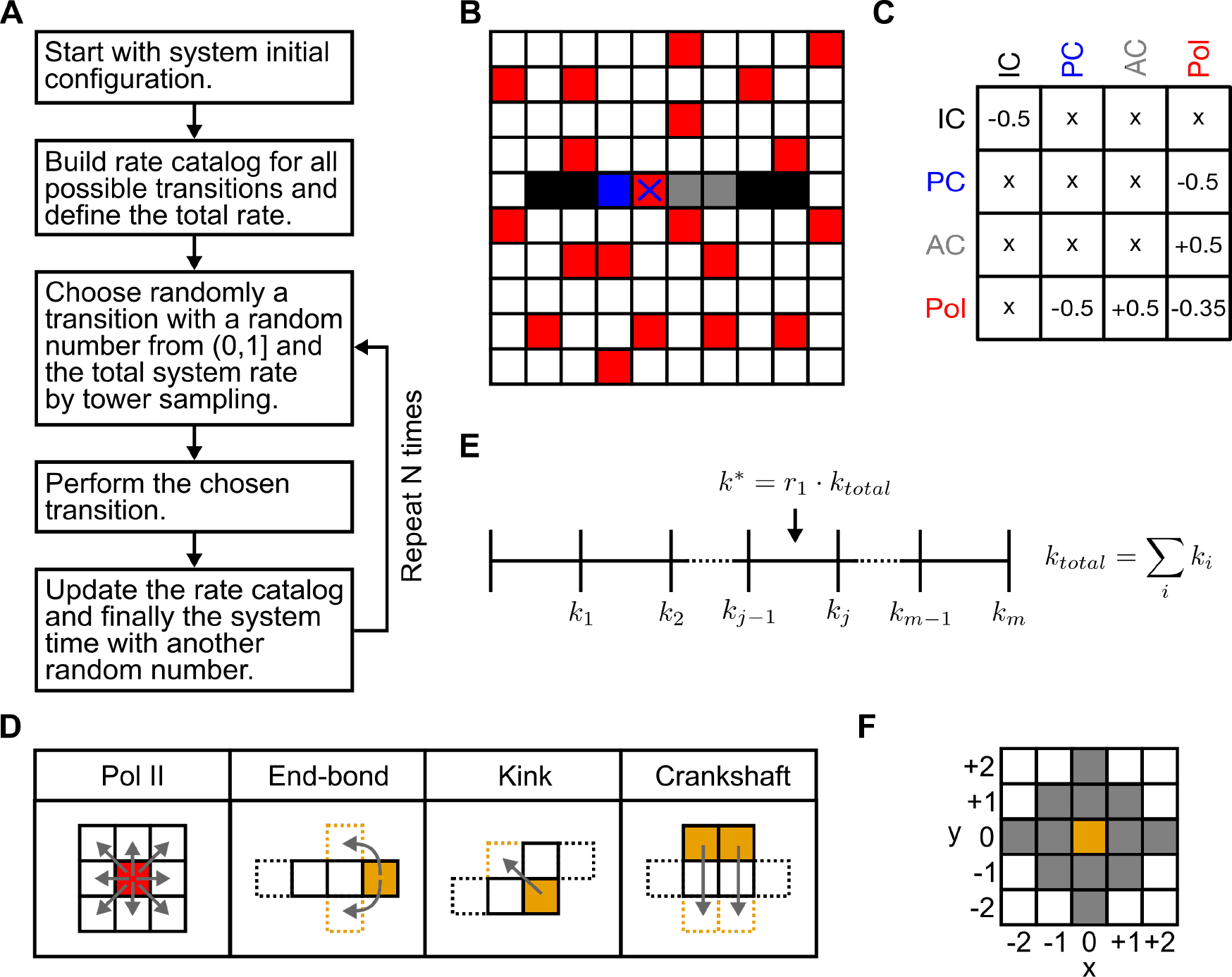
Lattice Kinetic Monte Carlo (LKMC) model summary. A) Algorithm overview. B) Illustration of a possible initial configuration containing a single polymer and all species within the system (black, blue, gray and red). C) Overview table of the interspecies affinity parameters. Table entries represent *w* values, crosses correspond to no interaction (*w* = 0). D) Possible move types for the different species. Pol II can move to all its eight nearest neighbours. Chains representing chromatin can undergo the Verdier-Stockmayer move set. E) Tower-sampling method to choose a transition. F) Minimal area for the local update of possible transitions.

**Fig. 11:**
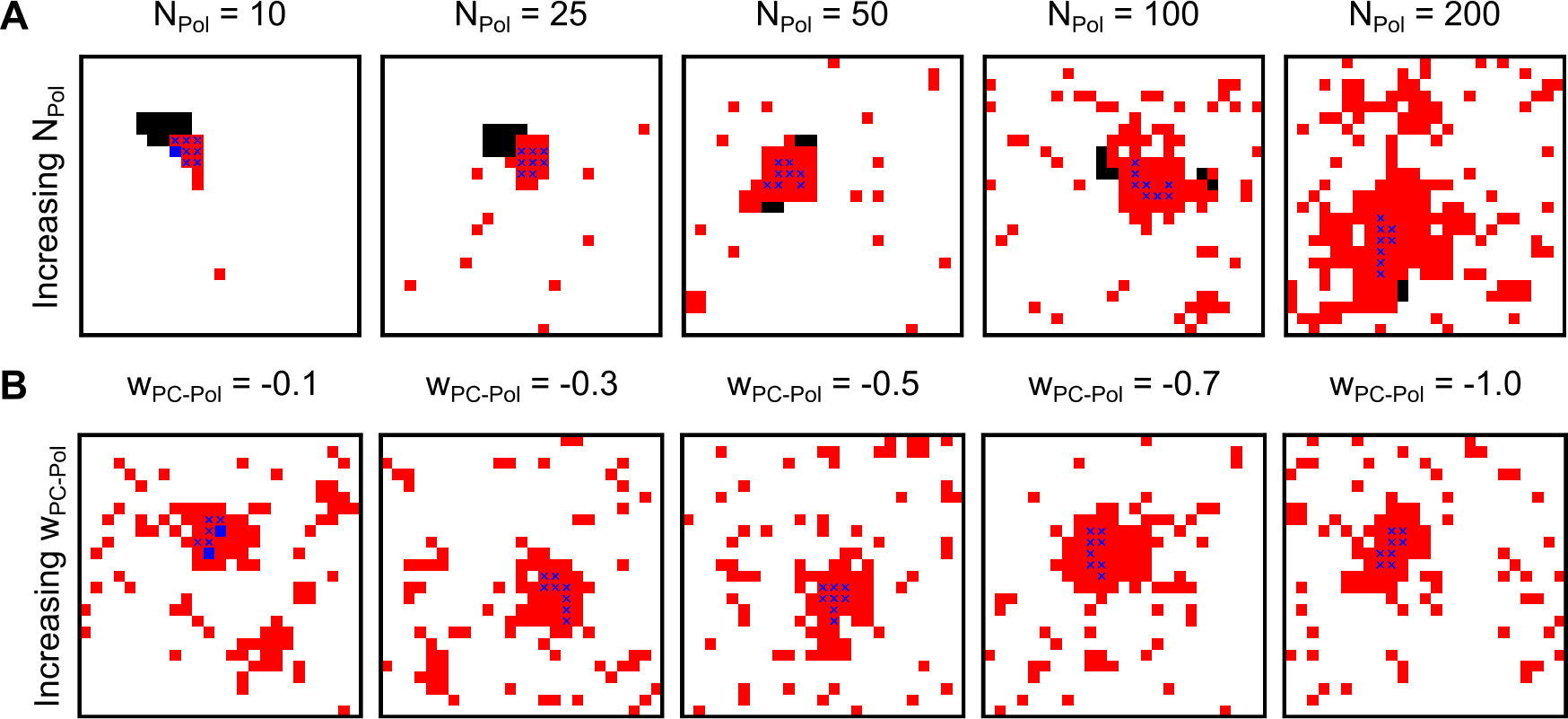
Parameter adjustment. A) Adjustment of *N_Pol_*. A single polymer chain of length *L_Polymer_* = 20, containing 8 blue particles was placed in the simulation. *w_Pol-Pol_* = −0.35 and *w_PC-Pol_* = −0.5 were held at constant values. A range of *N_Pol_* = 10,25,50,100,200 was used in different simulations as indicated. B) Adjustment of *w_PC-Pol_*. A single polymer chain of length *L_Polymer_* = 8 with only blue particles was placed in the simulation. *w_Pol-Pol_* = −0.35 and *N_Pol_* = 100 were held at constant values. A range of *w_PC-Pol_* = −0.1, – 0.3, – 0.5, – 0.7, – 1.0 was used in different simulations as indicated.

**Fig. 12:**
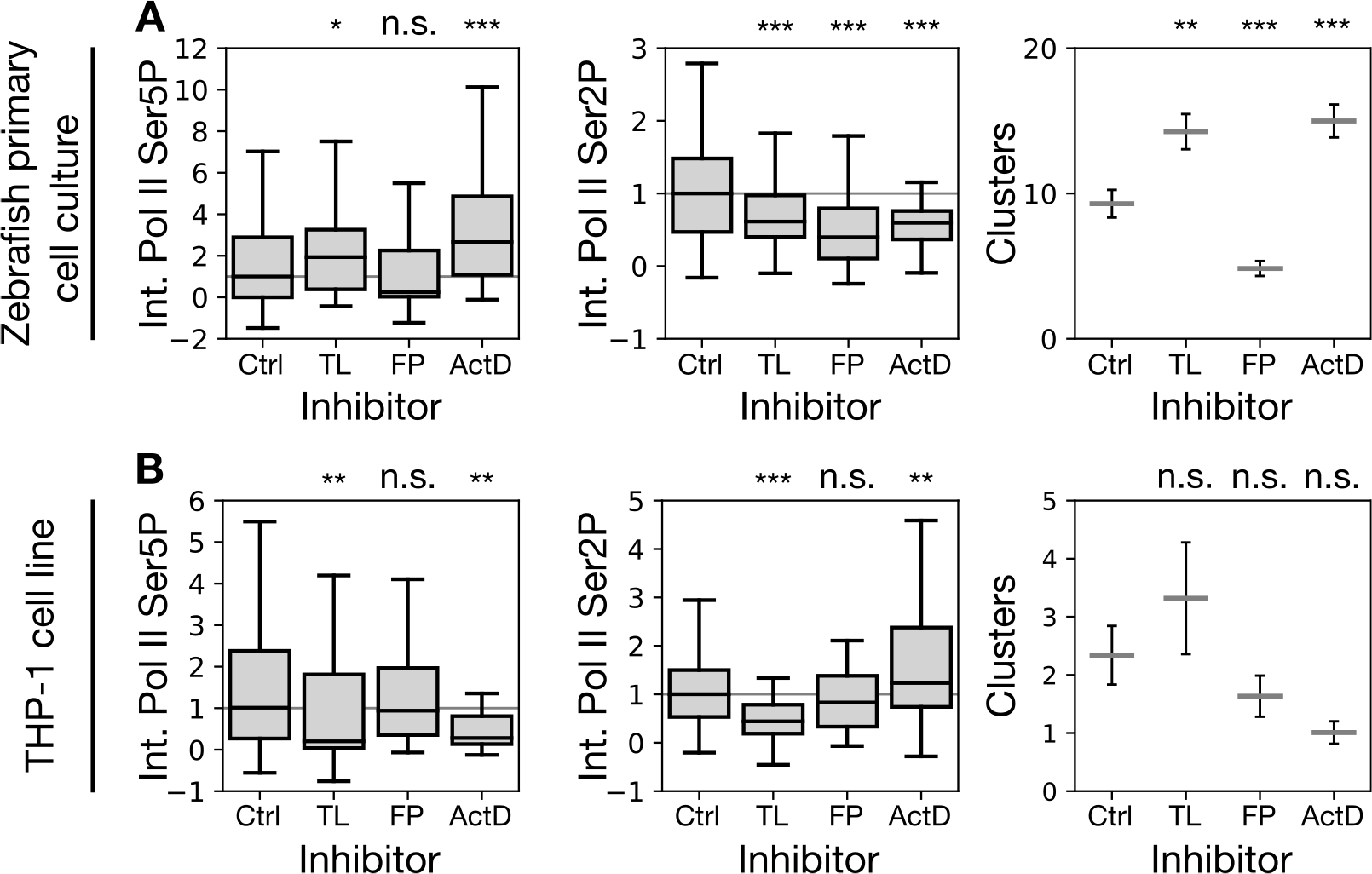
Inhibitor effects at whole nucleus level. A) RNA polymerase II Ser5 and Ser2 phosphorylation throughout the nucleus as obtained by immunofluorescence from primary cell cultures of zebrafish embryos, treated with the transcription inhibitors triptolide (TL), flavopiridol (FL), and actinomycin D (ActD). Intensity levels are shown as quantile plots (state quantiles), intensities are mean values after subtraction of cytoplasmic background, which can result in negative values. Within a given experimental repeat, values are normalized by the median of the control condition. Intensities are given as median with boxplots. The number of distinct clusters per nucleus is shown as mean±SEM. *** indicates *p* < 0.001, ** indicates *p* < 0.01, * indicates *p* < 0.05, n.s. indicates *p* ≥ 0.05 (Int. Pol II Ser5P: *p* = 0.04, *p* = 0.05, *p* < 0.0001, Int. Pol II Ser2P: *p* = 0.0002, *p* < 0.0001, *p* < 0.0001, Number of clusters: *p* = 0.004, *p* = 0.0002, *p* = 0.0006 for differences from the control condition, Bonferroni correction for multiple testing, *n* = 165,118,147,107 nuclei per condition from three independent experimental repeats.) B) Corresponding statistics for cultured THP-1 cells. (Int. Pol II Ser5P: *p* = 0.003, *p* = 0.61, *p* = 0.008, Int. Pol II Ser2P: *p* < 0.0001, *p* = 0.30, *p* = 0.009, Number of clusters: *p* = 0.10, *p* = 1.08, *p* = 0.10 for differences from the control condition, Bonferroni correction for multiple testing, *n* = 158,79,93,87 nuclei from three independent experimental repeats.)

**Fig. 13:**
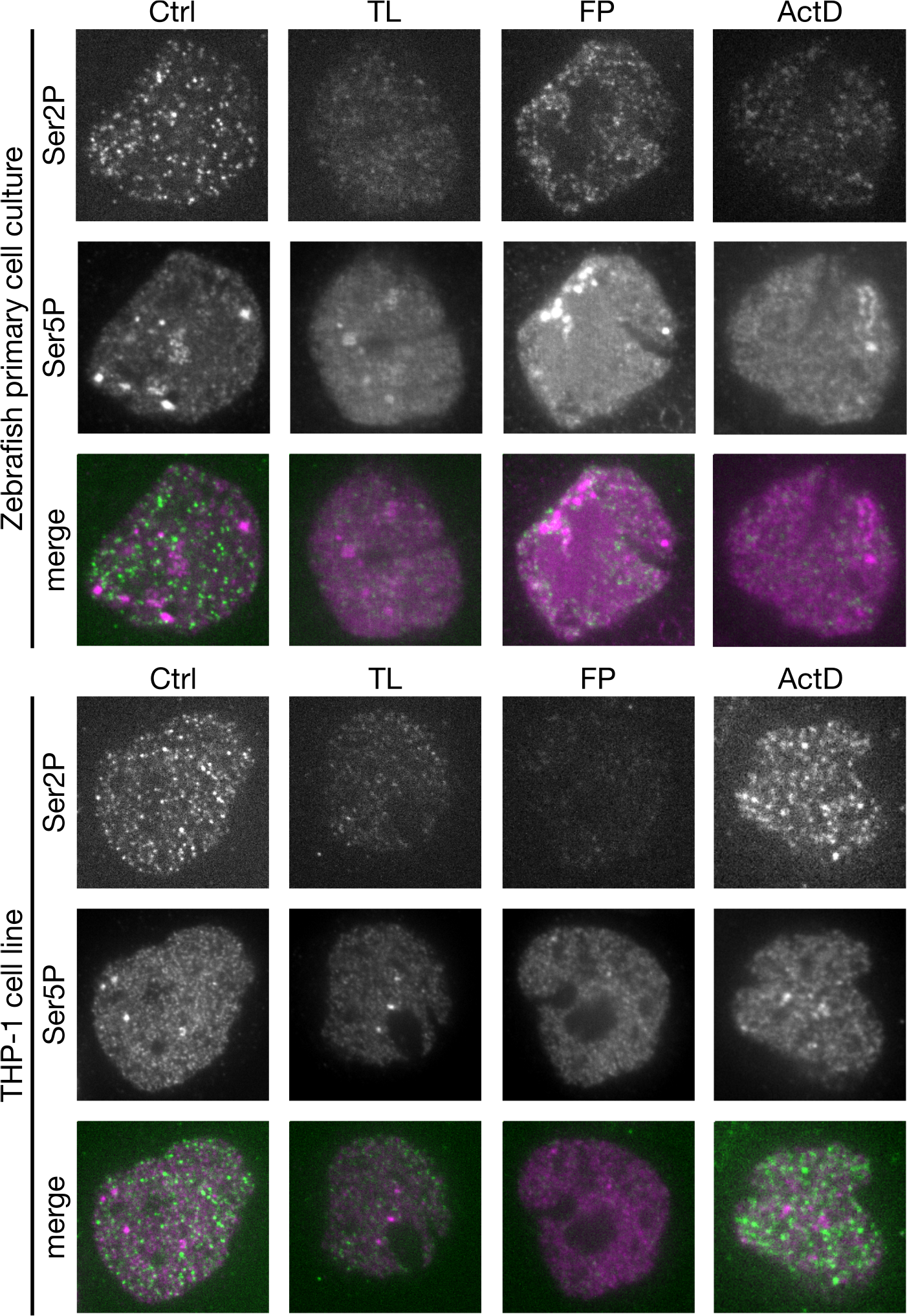
Sample images of clusters after inhibitor treatment. Representative mid-nuclear section nuclei in zebrafish primary cell culture (upper 3 rows) and THP-1 cell line (lower three rows). Cells were treated with control media (Ctrl), triptolide (TL), flavopidirol (FP) or actinomycin D for 30 min, and subsequently stained for Ser2 and Ser5 phosphorylation of RNA polymerase II by indirect immunofluorescence.

**Fig. 14:**
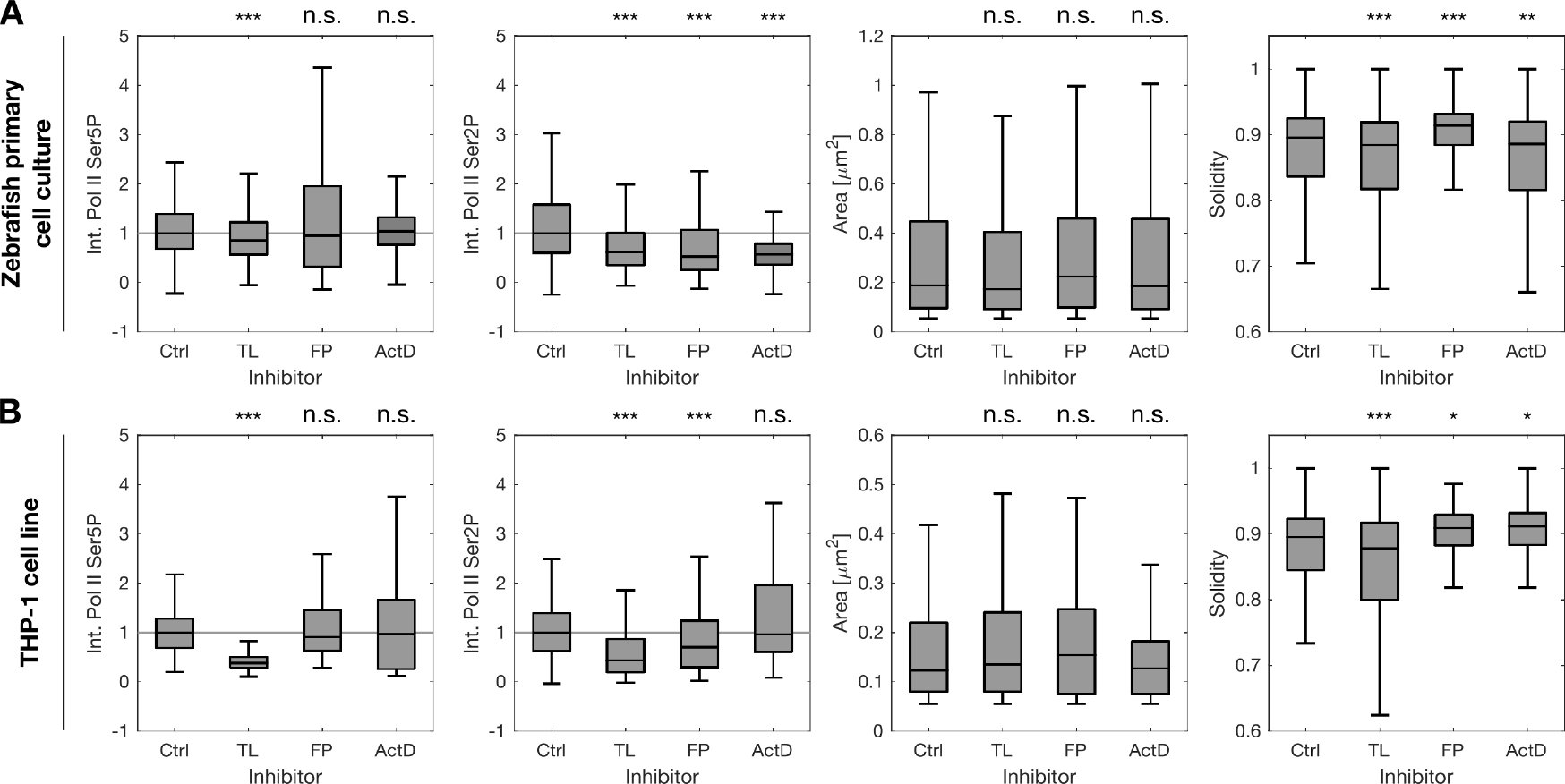
Changes in cluster properties upon inhibition. A) Pol II Ser5P and Pol II Ser2P mean intensity, area, and solidity of individual clusters in primary cell cultures established from oblong stage zebrafish embryos. Cultures were incubated for 30 minutes with inhibitors as indicated. Cluster intensities are mean intensities calculated over all pixels of maximum-projected clusters, cytoplasmic staining background was subtracted, and values were normalized by the median intensity of the control samples of a given experimental repeat (indicated by lines at intensity level 1). Shown are median values with boxplots. *** indicates *p* < 0.001, ** indicates *p* < 0.01, * indicates *p* < 0.05, n.s. indicates *p* ≥ 0.05 (Int. Pol II Ser5P: *p* < 0.0001, *p* = 0.25, *p* = 0.10, Int. Pol II Ser2P: *p* < 0.0001, *p* < 0.0001, *p* < 0.0001, Area: *p* = 0.26, *p* = 0.06, *p* = 2.74, Solidity: *p* = 0.0007, *p* < 0.0001, *p* = 0.001 for differences from the control condition, Bonferroni correction for multiple testing, *n* = 1514,1631,703,1567 clusters per condition from three independent experimental repeats.) B) Corresponding properties of clusters in undifferentiated THP-1 cells treated exactly like zebrafish primary cell culture. (Int. Pol II Ser5P: *p* < 0.0001, *p* = 0.36, *p* = 1.90, Int. Pol II Ser2P: *p* < 0.0001, *p* < 0.0001, *p* < 2.01, Area: *p* = 1.19, *p* = 0.10, *p* = 2.78, Solidity: *p* = 0.0004, *p* = 0.03, *p* = 0.01 for differences from the control condition, Bonferroni correction for multiple testing, *n* = 402,294,150,114 clusters per condition from three independent experimental repeats.)

**Fig. 15:**
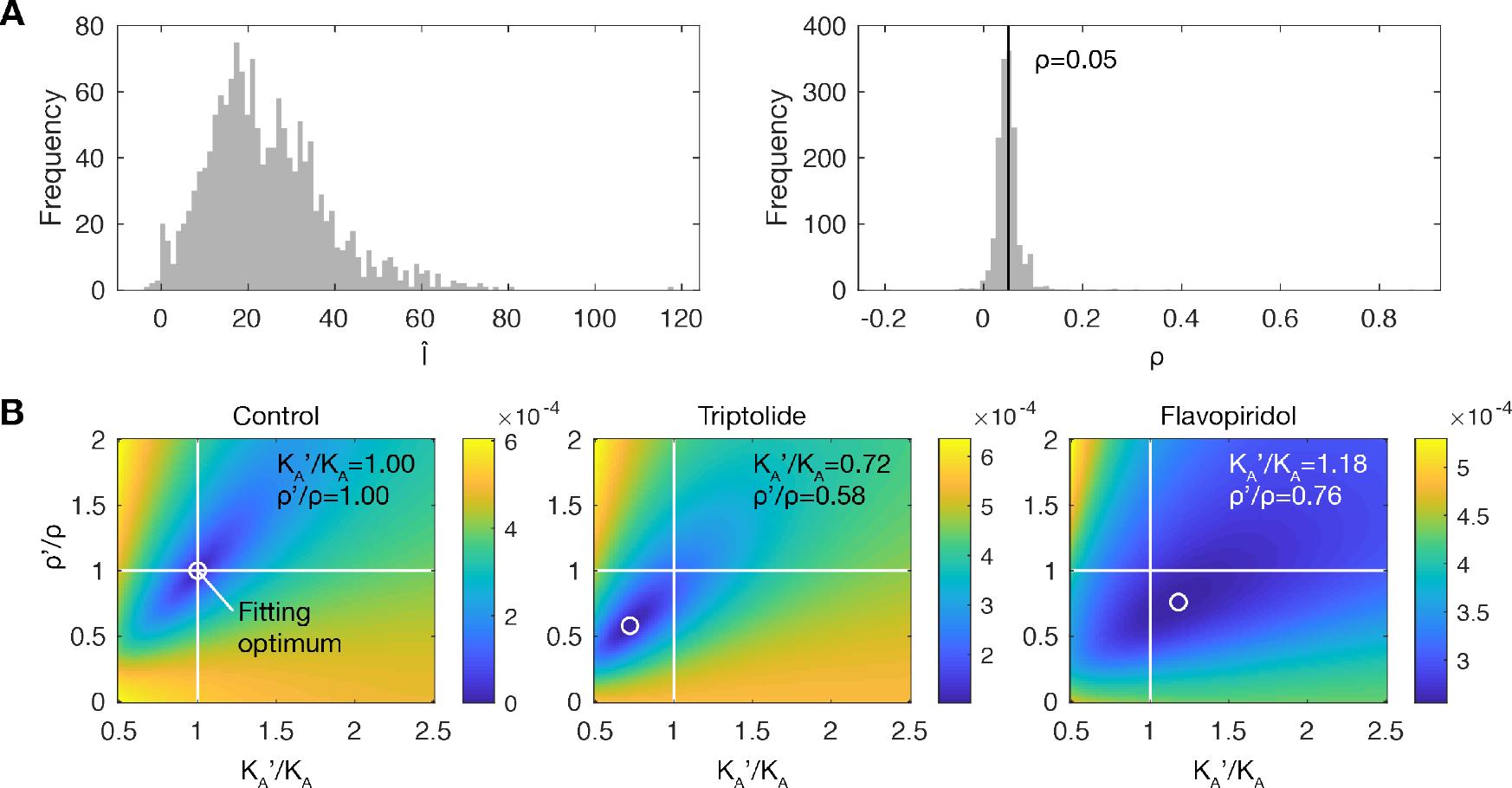
Fitting of model parameters to experimental data. A) Histograms of *I* and *p* values obtained using Pol II Ser5P and Pol II Ser2P intensity levels of Pol II clusters control-treated primary zebrafish embryo cell cultures. In the calculation of (*Î,ρ),* the parameter values *K_A_* =0.1 and *C* =1 were used. These were taken from previous work: *K_A_* = *k_A_/k_D_* = 0.1 considers *k_A_* ≈ 1 min^-1^ and *k_D_* ≈ 10 min^-1^^45^. *C = k_R_Telong* ≈ 1 considers *k_R_* ≈ 2 min^-1^^45^, length of 5 — 10 kb in zebrafish embryos at the late blastula stage of development^67^ and an elongation rate of 2 kb/s^68^. To assess the validity of this parameter calculation, the *ρ* histogram was compared against the expected *ρ* = *k_R_/k_D_* ≈ 0.05 (*k_D_* ≈ 1/7s,*k_R_* ≈ 1/150 s, previously estimated^45^, as indicated by a black line. This comparison suggests strong agreement. B) Fitting of the modified affinity values 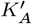 and *ρ*’ for cell cultures treated with DMSO (Control), triptolide, and flavopiridol. Images show the root-mean-square residuals, with white circles indicating the position of the minimal value. Residuals were calculated between the distributions of Pol II Ser5P and Pol II Ser2P intensities obtained from the treatment experiments directly or predicted based on the control condition and a modification of the *K_A_* and *ρ* values. The optimal values of 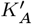 and 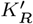 were chosen based on the location of the minimum.

## Notes

### Competing Interest Statement

The authors have declared no competing interest.

